# circAβ-a RNA encoded Aβ175—the hidden driver of β-amyloid plaque formation and deposition in sporadic Alzheimer’s disease

**DOI:** 10.1101/2025.01.18.633698

**Authors:** Dingding Mo, Yi Zhao, Yuxuan Liu, Juergen Brosius, Gelei Xiao

## Abstract

Mechanisms that trigger Aβ production in sporadic Alzheimer’s disease are still obscure. We recently reported the expression of a human circular RNA (circAβ-a) encoded Aβ peptide precursor variant (Aβ175). Presently, we demonstrated that AAV9 virus-expressed circAβ-a gave rise to extensive extracellular Aβ plaque depositions and microglial activation in mouse brain; this recapitulates critical pathogenic hallmarks within a sporadic AD mouse model. Specifically developed antibodies detected robust endogenous Aβ175 expression in HEK293 cells and hNSC-derived human neurons, underscoring the potential of Aβ175 as a salient Aβ precursor. Furthermore, we detected high levels of Aβ175 oligomers in young-adult human brains. In intermediate and old-age human brain samples, accumulation of soluble Aβ175 pentamers was reduced and Aβ175 oligomers were components of most insoluble Aβ plaques in older human brain. We propose a causal relationship between human circAβ-a RNA expression, dysregulation of Aβ175 oligomer processing/aggregation and Aβ plaque accumulation in sporadic AD.

## Introduction

Alzheimer’s disease (AD) is the most common cause of dementia; the detrimental neurodegenerative disorder results in the gradual loss of brain functions such as memory and cognition^1,2^. The key hallmarks of Alzheimer’s disease are the formation of extracellular β-amyloid (Aβ) plaques and intracellular neurofibrillary tangles (NFTs) in patient brains^2^. Although the actual role of Aβ in the pathogenesis of AD is still controversial^3,4^, extensive studies, including human genetics, have confirmed that Aβ peptide is closely associated with the development of AD pathologies^1,5,6^. Recent clinical trials of anti-Aβ immunotherapy on early-stage AD patients further support a significant causal role of Aβ neural toxicity^7^.

Previous studies established that Aβ can be produced from β-amyloid precursor proteins (APP) through sequential proteolysis by β-and γ-secretase^6,8^. Mutations in the protein coding sequence of APP and presenilin (PS1, PS2, the catalytic subunit of the γ-secretase complex) could accelerate the process and these mutations are associated with elevated Aβ levels and amyloid plaque deposition in the brain of familial AD patients^5,9–11^. However, gene mutations underlying familial AD only account for less than 1–6% of all AD cases globally^12^, and they fail to explain the sporadic AD patient majority lacking specific mutations in the *APP* and presenilin genes. Although β-secretase is the key enzyme that controls the rate-limiting step in the amyloidogenic pathway, it processes only very small amounts of the total human fl-APP^13^. βCTF (β-secretase cleaved product of APP) is a rare and transient metabolite in iPSC-derived human neurons^14,15^. On the other hand, the competitive non-amyloidogenic cleavage by α-secretase is predominant in human cells^16^. It has been shown that γ-secretase inhibition results in significant accumulation of α-secretase cleaved products (αCTF) in human HEK293 cells and iPSC-derived human neurons (Fig. 2A in^17^ and Fig. 5G in^14^). Taken together, previous studies strongly imply that fl-APP may not be the main source of Aβ peptides. This points to hitherto undiscovered mechanism(s) underlying the accumulation of Aβ in brains of sporadic AD patients.

Circular RNA (circRNA) is formed during processing of primary transcripts^18^. Most circular RNAs are derived from (parts of) primary transcripts transcribed from protein-coding genes via back-splicing, and, reportedly, are involved in various biological processes^18^. In the brain, circular RNAs are particularly abundant, suggesting neural-related functions^18^. Certain circRNAs have the potential to be translated into polypeptides^19–22^. We recently found that the APP gene can also generate a circular RNA (circAβ-a, hsa_circ_0007556) encoding a precursor (Aβ-related protein, Aβ175) of the Aβ peptide^23^. Translation of circAβ-a elongates beyond the junction of the circle, thus leading into a different reading frame for 17 amino acids until a translational stop codon is encountered^23^. Consequently, Aβ175 differs with respect to its C-terminus from the canonical fl-APP-derived CTFs. The Aβ peptide is thought to be the key pathological factor of Alzheimer’s disease. Our data indicate that the dominant pathogenic potential may lie within the circAβ-a encoded peptides^23^. Since the biogenesis and translation of circAβ-a does not appear to require APP gene mutations, the neurotoxicity of its translation product is a potential explanation for the development of Alzheimer’s disease in patients devoid of familial mutations^23^.

Alzheimer research lacks suitable animal models that represent the origins and development of pathological features of sporadic Alzheimer’s disease, thus limiting the efforts to transform genetic knowledge into clinical applications^24^. Previous animal models of familial Alzheimer’s disease relied on double-transgenic or even triple-transgenic mice that overexpress mutated human APP, PSEN1, and TAU proteins to obtain amyloid plaques and tangles in the mouse brain^24^. The APP knock-in mouse model of the humanized Aβ sequence and Swedish mutations in exon 16, Beyreuther/Iberian and Arctic mutations in exon 17, showed classic Aβ pathology, neuroinflammation and some degree of memory impairment in an ageing-dependent manner^25^. However, all these models are far from the actual pathology of patients and there is no good clinical recapitulation of human AD pathogenesis. As described previously, extensive analysis of fl-APP protein expression in the brains of patients showed that APP is neither mutated nor overexpressed in most cases. Therefore, the overexpression of mutated human fl-APP protein required to produce Aβ in mouse brains may cause *in vivo* artifacts^26^. Therefore, the obvious shortcomings and limitations of currently available mouse models render them unreliable to correctly evaluate the clinical effects of drug candidates in humans^26^.

Overexpression of wild type human fl-APP protein in mouse brain does not produce significant amounts of Aβ, and there is no Aβ plaque formation in these animals^26–28^. Due to the fact that the implemented transgenes are intron-less cDNAs, no human circAβ-a RNA is produced. In contrast, overexpression of wild type human circAβ-a RNA in the mouse brain may reflect more closely the nature of sporadic AD, as the translated product of circAβ-a RNA itself may have the ability to produce enough Aβ and generate extracellular Aβ plaques without any mutations; this would provide a useful animal model for drug screening. To achieve such a goal, we constructed an AAV9 virus harboring the circAβ-a RNA expression cassette^23^ under the human neuron-specific SYN1 promoter (AAV9-circAβ-a). After injection of the AAV9-circAβ-a into the frontal cortex region of the mouse brain, we observed robust Aβ accumulation and plaque distributions, using a GFP expressing virus as control.

Furthermore, we developed specific antibodies in mice against the unique C-terminus of the human circAβ-a translation product, Aβ175. Western blot analysis showed robust Aβ175 expression in human HEK293/HEK293T cell lines, hNSC-derived human neurons and human brains, supporting its potential role as the key source of Aβ in humans. Already, in the brains of ∼20-year-old individuals, we observed a robust presence of Aβ175 oligomers; dysregulation, perhaps at the processing level, may begin at age ∼30 being augmented further with advancing age. Significantly, we found that Aβ175 aggregated into the insoluble Aβ plaques in older human brains. Notably, fl-APP is predominately processed by α-secretase to generate non-amyloidogenic products and knock-down of Aβ175 and knockout of fl-APP protein in HEK293 cells indicated that Aβ175 was the main precursor to Aβ peptides, but not fl-APP.

These multiple lines of evidence point to the dysregulation of Aβ175 oligomer processing and their accelerated aggregation during ageing as the major origin of Aβ plaque accumulations in sporadic AD. The specific antisense oligonucleotide (ASO) targeting the circAβ-a RNA could significantly reduce endogenous Aβ175 expression and the antibodies targeting the Aβ175 component of Aβ plaques may serve as the basis for developing disease-modifying agents in sporadic AD treatment.

## STAR Methods

### AAV9-circAβ-a plasmid construction, AAV9 virus preparation and injection

The circAβ-a RNA (Human hsa_circ_0007556 (circBase^29^)) was used in the expression cassette (with intron mediated enhancement^19^), amplified from pCircRNA-DMo-Aβ-a^23^ and inserted into an AAV2 backbone vector under the human SYN1 promoter. The construct was sequenced and transfected into HEK293 cells for AAV9 virus production (designated as AAV9-circAβ-a) using standard protocol. The virus titer was calculated with qPCR (3.48 × 10^13^ GC/ml).

For a single virus injection, 0.70 × 10^10^ GC or 6.26 × 10^10^ GC AAV9-circAβ-a was injected into layer V of the left frontal cortex region in nine-month-old male C57BL/6 mice. As a negative control, we injected into the same brain region with 0.70 × 10^10^ GC of AAV9-Gcamp7f virus, which expresses cpGFP, a green fluorescent protein (GFP) variant, under the human SYN1 promoter. For some mice, we co-injected the same brain region with both AAV9-circAβ-a and AAV9-Gcamp7f virus.

### RNA isolation and reverse transcription-polymerase chain reaction (RT-PCR)

Total RNA isolation from the AAV9 virus injected region of the left frontal cortex or from HEK293/HEK293T cells was performed as previously described^19^ using TRIzol LS Reagent (10296010, Invitrogen). For cDNA synthesis, 0.5 µg of total RNA was used as template for reverse transcription with the SuperScript III First-Strand Synthesis System (18080044, Invitrogen) and random hexamer primers. PCR was performed with PrimeSTAR® Max DNA Polymerase (R045B, TaKaRa) and previously established circAβ-a primers (circAβ-a-R1 and circAβ-a-F1)^23^ for 40 cycles. RT-PCR of circAβ-a was performed as previously described^23^.

### Immunofluorescence (IF) of brain tissue sections

Mouse brain slices were collected three, four or seven months after AAV9-circAβ-a injection to the left frontal cortex of nine-month-old male mice, using the following procedure: anesthetized mice were perfused with PBS and 4% paraformaldehyde (PFA). Mouse or human brain samples were fixed with 4% PFA for 24 hours followed by a 24-hour dehydration initially in 15% sucrose and another 24-hour dehydration in 30% sucrose. Sections (30–50 μm) were generated in a Leica CM1950 cryostat. Mounted sections were treated with 70% formic acid for 4 minutes or heated in citrate buffer (10 mM trisodium citrate, 0.05% Tween 20, pH 6.0) at 95 °C for 20 minutes and thrice washed with PBS. 4G8 antibody (800708, BioLegend) against human Aβ was used in immunohistostaining incubating at 4 °C overnight, followed by incubation with Alexa Fluor® 546 goat anti-mouse IgG secondary antibody (A-11003, Invitrogen) for one hour at room temperature. Anti-GFP (chicken) and goat anti-chicken IgY (H+L), Alexa Fluor™ 488 (A-11039) were used for cpGFP immunohistostaining. The immunofluorescence analysis of Aβ175 oligomers in human brain was performed with mouse monoclonal antibody against Aβ175 oligomers (MO-05-158) and Alexa Fluor® 546 goat anti-mouse IgG secondary antibody (A-11003, Invitrogen). Human brain sections were also incubated with β-amyloid (1-40) rabbit mAb (A24949, ABclonal) and Alexa Fluor™ 633 goat anti-rabbit IgG (H+L) cross-adsorbed secondary antibody (A-21070, Invitrogen). For co-staining of Aβ plaques, 1 µM thioflavin-S (in PBS) (T1892, Sigma) or 1 µM QM-FN-SO_3_ (in PBS) (gift from Prof. Zhiqian Guo, East China University Science and Technology) was added to the brain sections for one-hour incubation at room temperature and washed with PBS four times. Sections were also stained with X-34 staining solution (10 µM X-34 (SML1954, Sigma), 60% PBS (vol/vol), 40% ethanol, 20 mM NaOH) for 20 minutes at room temperature. DAPI Fluoromount-G® (0100-20, SouthernBiotech) or Fluoromount-G® (0100-01, SouthernBiotech) mounting medium was used to seal the slide coverslips. Images were generated with a ZEISS LSM 900/980 Confocal Microscopy System.

### HEK293/HEK293T cell culture and ASO transfection

pCMV-Aβ175 was transfected into HEK293T as described above to express the Aβ175 protein as positive control^23^. For antisense oligonucleotide (ASO) inhibition of circAβ-a RNA, 300 or 100-500 nM antisense oligonucleotide (circAβ-a-ASO) targeted at the junction region of circAβ-a RNA was transfected into HEK293/HEK293T cells with Oligofectamine transfection reagent (12252011, Invitrogen). After 2 days of culture in DMEM medium with 1% FBS, cells were collected for total RNA isolation using TRIzol reagent (Ambion) and total protein was prepared with RIPA buffer (R0278, Sigma). A scrambled ASO was used as negative control. circAβ-a-ASO: 5’ mA*mA*dGdCdAdGdCdTdCdAdTdCdTdCdC dAdCdCdAdC*mA*mC 3’; scrambled control: 5’ dC*dT*dCdCdAdGdCdTdAdCdA dCdAdAdGdCdAdCdCdC*dA*dT 3’; *, phosphorothioate modification sites. mA, 2’-O-methyl-adenosine; mC, 2’-O-methyl-cytosine.

### CRISPR-Cas9 genome editing of APP exon in HEK293 cells

The guide RNA sequence of CRISPR was designed against exon 13 of the human APP gene (5’ TGAATCTCCTCGGCCACTGC 3’). The guide RNA sequence was cloned into the genome editing plasmid described^30^: pSpCas9(BB)-2A-Puro (PX459) V2.0 was a gift from Feng Zhang (Addgene plasmid # 62988; http://n2t.net/addgene:62988; RRID: Addgene_62988). HEK293 cells were transfected with the genome editing plasmid and selected with puromycin. Cell lines with edited APP exon13 were established. The cells were further verified by Western blot with KO validated APP rabbit mAb (A17911, ABclonal) and Aβ175 antibody (MO-05).

### shRNA plasmid and shRNA lentivirus

circAβ-a shRNA plasmid was constructed under the U6 snRNA promoter by standard methods. shRNA sense sequence against circAβ-a junction region was “5’ GTGTGG TGGAGATGAGCTGCT 3”. The shRNA knock-down lentivirus was produced with a standard protocol in HEK293T cells with MISSION^®^pLKO.1-puro vector. The infected HEK293 cells were selected with puromycin. The established cell lines were verified by Aβ175 Western blot employing MO-05 antibody. The sense sequence of shRNA against the second exon of human APP mRNA was 5’ CATGCACATGAAT GTCCAG 3’ as previously described^31^. The shRNA sequence was inserted into the MISSION^®^pLKO.1-puro vector. The lentivirus was prepared similarly.

### Human neuron culture

A human neural stem cell (NSC)-derived neuron culture was established as described^32^. The shRNA knock-down lentivirus infected human neural progenitor cells (NPCs) were selected with puromycin. After 14 days of neural differentiation, human neuron cultures were collected for Western blot analysis.

### Antibody production

The antigen (derived from peptide CFRKSKTIQMTSWPT) was injected into mice using standard protocols. Positive polyclonal antibody was selected and named as MO-05. The selected immune cells were hybridized with immortal Sp2/0 cells. The supernatants of established hybridized cell lines were tested with Western blot analysis of Aβ175 monomers and oligomers. Positive clones were selected and four monoclonal antibodies were purified and designated as MO-05-158, MO-05-721, MO-05-160, MO-05-165. In addition, MO-05-158 and MO-05-165 were humanized by antibody engineering using a standard strategy and designated as humanized MO-05-158 and humanized MO-05-158, respectively. Humanized antibodies were expressed in HEK293 cells by transient transfection and purified by affinity chromatography using standard protocols.

### Aβ indirect enzyme-linked immunosorbent assay (ELISA) measurement

Aβ peptide presence in the media of these established HEK293 cell lines (fl-APP-KO and circAβ-a shRNA knock-down) were analyzed by indirect Aβ ELISA measurement with the 0.5 μg/mL purified 4G8 monoclonal antibody (800712, BioLegend) (1:2000 dilution).

### Western blot

Western blots were performed as previously described^23^. Briefly, HEK293, HEK293T cells, human hNSC-derived neurons, brain cortex tissue from mouse or human (detailed information is provided in Supplementary Table 1) were prepared in RIAPA buffer with protease and phosphatase inhibitor. Forty µg total protein extract was fractioned on 8–20% precast protein gels and transferred to 0.2 µm pore-size nitrocellulose membranes (66485, Pall). For monomer Aβ detection, the pellets of brain tissue lysed in RIAPA buffer were extracted with 70% formic acid and neutralized with 5 N NaOH in 1 M Tris solution. The resulting solutions were fractioned on 8–20% precast polyacrylamide gels and transferred to 0.2 µm pore-size PVDF membranes (ISEQ00010, Millipore). Blot membranes were heated for 5 minutes in PBS to retrieve epitopes prior blocking. 4G8 monoclonal antibody (800708, BioLegend) was used in Aβ immunoblotting. Purified 4G8 monoclonal antibody (800712, BioLegend) was used in βCTF175 trimer and Aβ175 blotting. Anti GFP-Tag mAb (AE012, ABclonal) was used for Gcamp7f detection. Anti-GAPDH (AC002, ABclonal) or β-actin (A5441, Sigma) were used as loading control. Aβ175 was detected by a newly developed polyclonal antibody against its unique C-terminal sequence (MO-05) in mice. The monoclonal antibodies (MO-05-158, MO-05-160, MO-05-165 and MO-05-721 produced in this study) were used to detect Aβ175 oligomers. APP full length protein (fl-APP) was detected by KO validated APP rabbit mAb (A17911, ABclonal). For Western blots of human neuron cultures, βIII-tubulin rabbit mAb (A17913, ABclonal) was used as a neuron-specific marker. HRP goat anti-mouse IgG (H+L) (AS003, ABclonal) and HRP goat anti-rabbit IgG (H+L) (AS014, ABclonal) were used as secondary antibodies. Signals were developed with Super ECL Plus solution (S6009M, UElandy Inc.) and visualized with the ChemiDoc MP Imaging System (Bio-Rad).

## Results

### *In vivo* expression of circAβ-a RNA in the mouse brain via AAV9 virus construct injection

To achieve circAβ-a expression in the mouse brain, we injected the AAV9 virus encoding the circAβ-a RNA expression cassette^23^ under the neuron-specific human SYN1 promoter (AAV9-circAβ-a, Fig. 1A), into the left frontal cortex of nine-month-old C57BL/6 mice. As an injection control, an AAV9 virus encoding GCaMP7f under human SYN1 promoter (AAV9-GCaMP7f), which expresses a cpGFP protein under the SYN1 promoter was co-injected. Mice were sacrificed 1.5 months post injection, and the injected brain regions were collected. Total RNA was isolated for RT-PCR to amplify the circAβ-a RNA with our previously established divergent primers^23^ (amplicon size: 499 bp) (Fig. 1B). After agarose gel electrophoresis, an expected RT-PCR product at the 500 bp ladder position was observed (Fig. 1B), indicating the generation of circAβ-a RNA. Furthermore, DNA sequencing of the RT-PCR products confirmed the human circAβ-a amplicon sequence (data not shown). Thus, human circAβ-a RNA was successfully overexpressed in mouse brain *in vivo*. The results are consistent with our previously established circAβ-a RNA expression in HEK293 cells^23^.

**Fig. 1.**
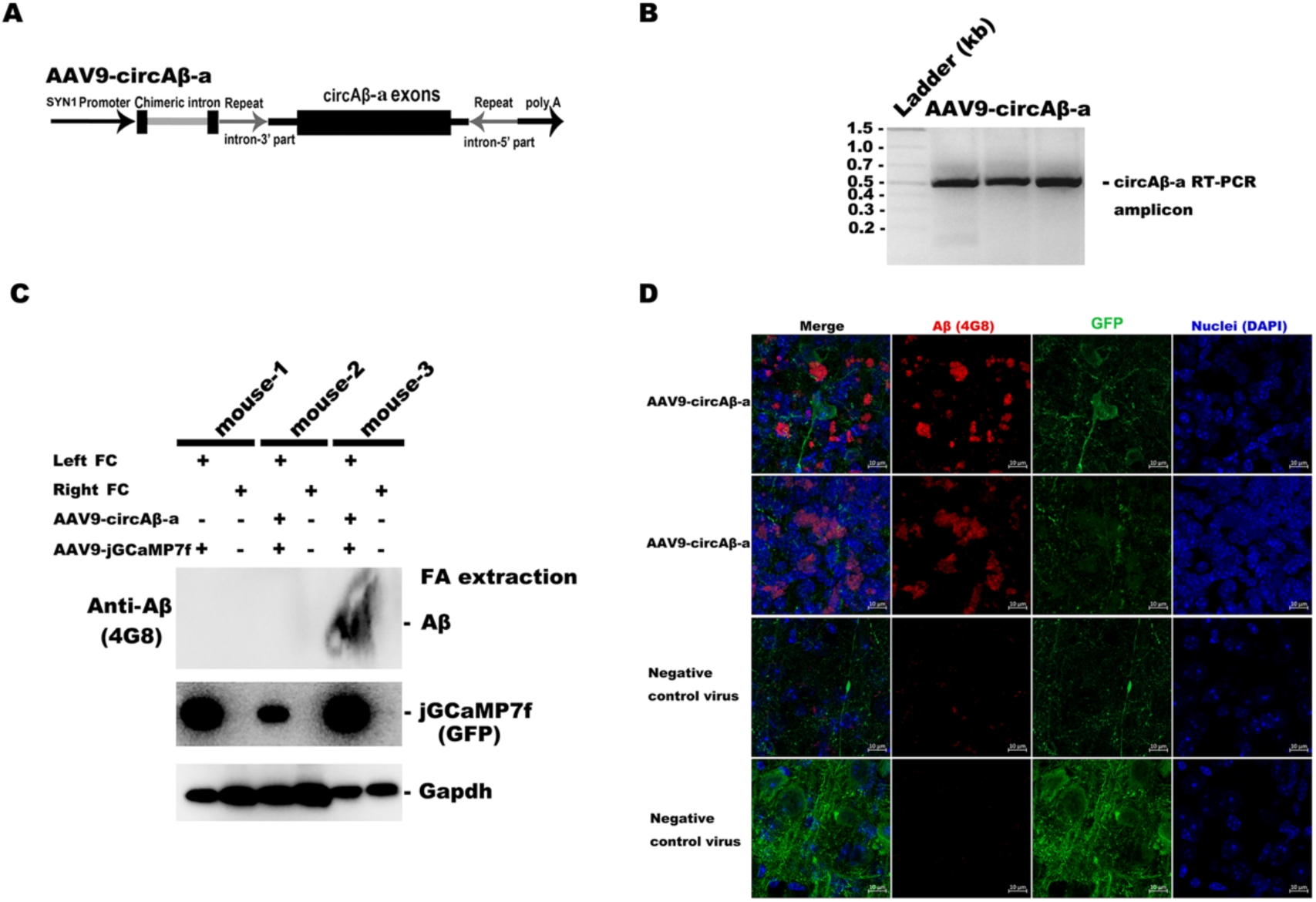
human circAβ-a RNA, Aβ expressions and Aβ plaque depositions in AAV9-circAβ-a injected mouse brains. A. Schematic diagram of the AAV9 virus expressing circAβ-a; B. RT-PCR of circAβ-a isolated from the frontal cortex of AAV9-circAβ-a injected mice; the amplicon is 499 bp; three mouse samples were used (N=3). C. Aβ expression in the virus injected mouse brains; Left FC, left frontal cortex; Right FC, right frontal cortex; Aβ were immunoblotted with 4G8 antibody; -, without virus injection; +, with virus injection. D. Aβ plaque immunostaining of frontal cortex after AAV9-circAβ-a injection; blue, DAPI (nuclei); red, Aβ (4G8); green, GFP; negative control virus, AAV9-Gcamp7f. Three mice were used in the experiments (N=3) and multiple slice images were captured; representative images were shown in the respective rows.

### Aβ expression from human circAβ-a RNA in virus injected mouse brain *in vivo*

As demonstrated in HEK293 cells, circAβ-a can act as a template for Aβ175 translation and Aβ peptides can be generated^23^. One and a half months post AAV9-circAβ-a injection into the brain of nine-month-old male C57BL/6 mice, total protein was extracted from the AAV virus injected region with RIPA buffer and Western blotting was performed with GAPDH and GFP antibodies as loading controls.

Undissolved pellets were further extracted with formic acid as detailed in Materials and Methods. These extracts were also used for Western blot (4G8) detection of Aβ. As shown in Fig. 1C, the left frontal cortex of mouse-1 injected only with AAV9-GCaMP7f did not reveal the corresponding signals, demonstrating that analogous injection of GFP marker virus did not affect/deregulate Aβ expression from the endogenous mouse *App* gene. Western blot of formic acid extracted pellets from the left frontal cortex of mouse-3 (6.26 × 10^10^ GC of AAV9-circAβ-a virus injected) revealed a heavy Aβ monomer band, yet the left frontal cortex of negative control mouse-1 failed to produce a signal (Fig. 1C). We did not detect Aβ monomers in formic acid extracted pellets of the left frontal cortex of mouse-2, presumably caused by the relatively lower injection titer (0.70 × 10^10^ GC) of AAV9-circAβ-a virus and the technical difficulty of low level Aβ monomer isolation and detection (as the GFP signal is relatively low in mouse-2, Fig. 1C). As an endogenous negative control, the un-injected right frontal cortex of mouse-3 was devoid of signal in the corresponding tissue areas (Fig. 1C). In conclusion, Western blots revealed Aβ peptides in the frontal cortex tissue of circAβ-a RNA overexpressed mouse brains.

### *In vivo* Aβ plaque depositions in the circAβ-a RNA expressing mouse brains

We examined the *in vivo* Aβ plaque formation capability of these Aβ peptides in mouse brain by using 4G8 antibodies. Extracellular Aβ plaques were identified in the brain slices of C57BL/6 mice four months after the injection of AAV9-circAβ-a virus into the nine-month-old males. As shown in Fig. 1D, the morphologies of 4G8 positive signals (red) in circAβ-a expressing tissue resembles the cotton wool Aβ plaques commonly found in AD patients^33^. A negative control showed very limited 4G8 positive signals in slices from AAV9-jGCaMP7f injected mice. As co-injection control, the green GFP signal was prominent in both circAβ-a expression and negative control mice.

Furthermore, we co-stained mouse brain slices with Aβ antibody (4G8) and plaque-binding chemical (Thioflavin S, Thio-S), the latter binding to the β-pleated sheet conformation in amyloid plaques. In the circAβ-a expressing AAV9 virus infected mouse brain slices, we detected significant Aβ plaque signals with both 4G8 antibody and Thio-S. Such co-localization further validated the identities of Aβ plaques (Fig.2A). Two additional Aβ plaque-detecting chemicals, namely QM-FN-SO_3_^34^ and X-34 were used to perform co-imaging of Aβ plaques with 4G8 antibody (Fig.2B, C). Once more, we obtained co-localized Aβ plaques by both 4G8 and QM-FN-SO_3_/X-34 (Fig.2B, C). Taken together, the co-staining with three plaque-specific chemicals and the specific Aβ antibody (4G8), solidly validated robust Aβ plaque formations and depositions in human circAβ-a expressing mouse brains (Fig.1D; Fig.2A, B, C).

**Fig. 2.**
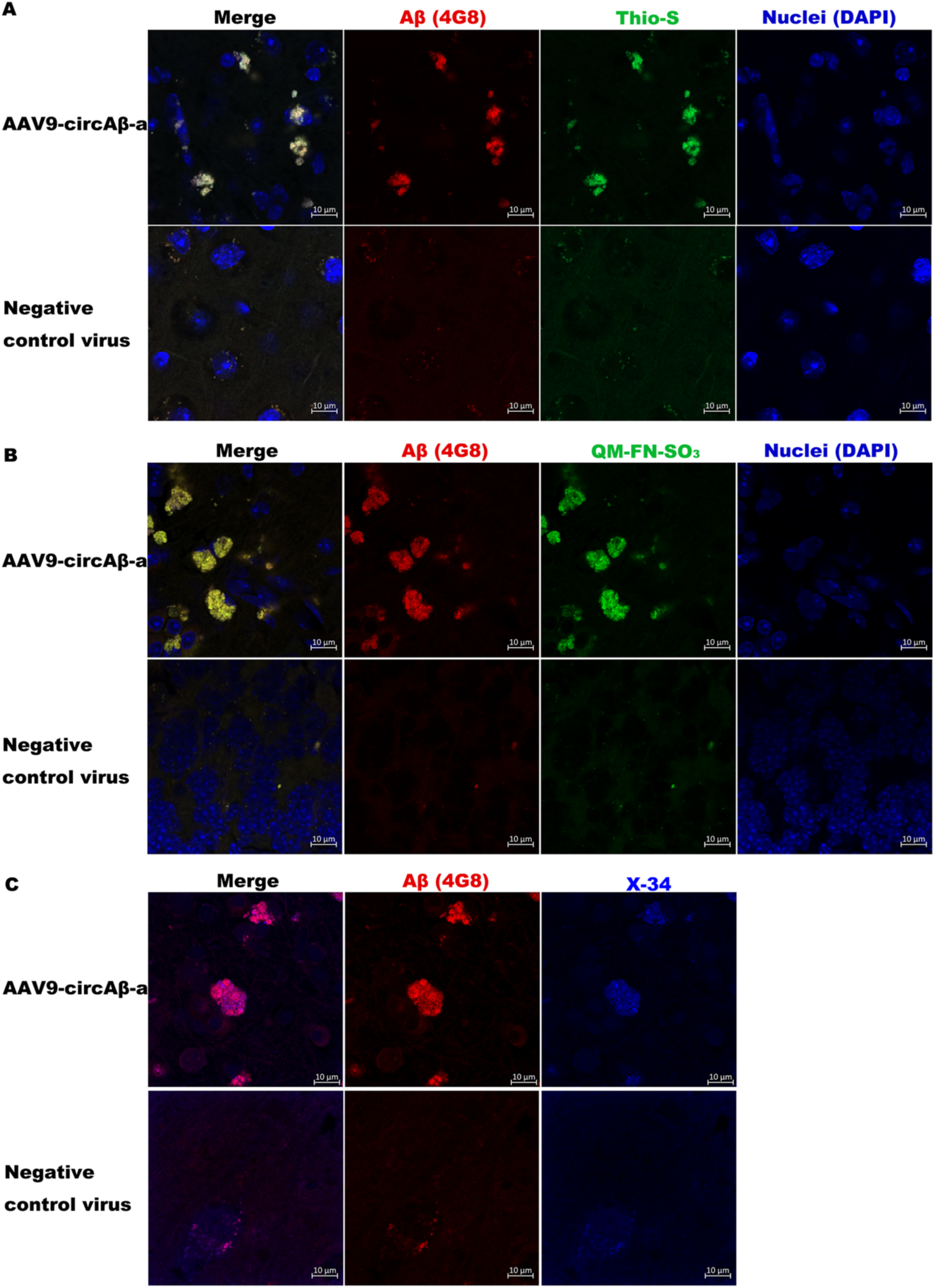
Co-staining of Aβ plaques in human circAβ-a RNA expressed mouse brain slices. **A**. blue, DAPI; green, Thioflavin S (Thio-S); red, Aβ (4G8); yellow, merged red and green. **B.** blue, DAPI; green, QM-FN-SO_3_; red, Aβ (4G8); yellow, merged red and green. **C.** red, Aβ (4G8); blue, X-34; Negative control virus, AAV9-Gcamp7f. At least three mice from each group were used in the experiments (N=3); multiple slice images were captured, and representative images were shown here.

These positive plaques were mostly located separately from the DAPI stained nuclei (Fig.1D; Fig.2A, B). Such extracellular deposits are one of the key characteristics of Aβ plaque depositions in human AD patient brains. Notably, the diameter size of these plaques was around 10 µm, relatively small compared to the average size of plaques in familial AD mouse models and AD patient brains^35,36^. Such differences may be due to the relatively small amount of AAV virus used for infection (6.26 × 10^10^ GC). Presumably, circAβ-a expressing transgenic mice will exhibit significantly larger Aβ plaques and Aβ plaques may grow larger with additional time (months) after virus injection. In fact, mice of the three different time points post AAV9-circAβ-a virus injection (3, 4 and 7 months) exhibited increasing Aβ plaques sizes (Supplementary Fig. 1), demonstrating not only the reproducibility of the *in vivo* experiments but also indicating the age-dependence of Aβ plaque growth in the human circAβ-a RNA expressing mouse brains.

### Microglial activation in the human circAβ-a RNA expressing mouse brains *in vivo*

Aβ plaque accumulation in human brains normally activates microglia, the immune cells that respond to the toxic protein aggregations in brains. To verify such a process in the human circAβ-a RNA expressing mouse brain, we incubated the brain slices with IBA1 antibody, which is a microglial activation marker. As shown in Fig. 3, around the green Thio-S positive region (Aβ plaques), there were significant IBA1 (red) signals in the AAV9-circAβ-a virus injected mouse, while the negative control virus injection exhibited neither green Thio-S, nor strong red IBA1 signals. This indicated that accumulation of Aβ plaques activated microglia in the circAβ-a RNA expressed mouse brains, paralleling the microglial activation of Aβ pathology in human AD brains.

**Fig. 3.**
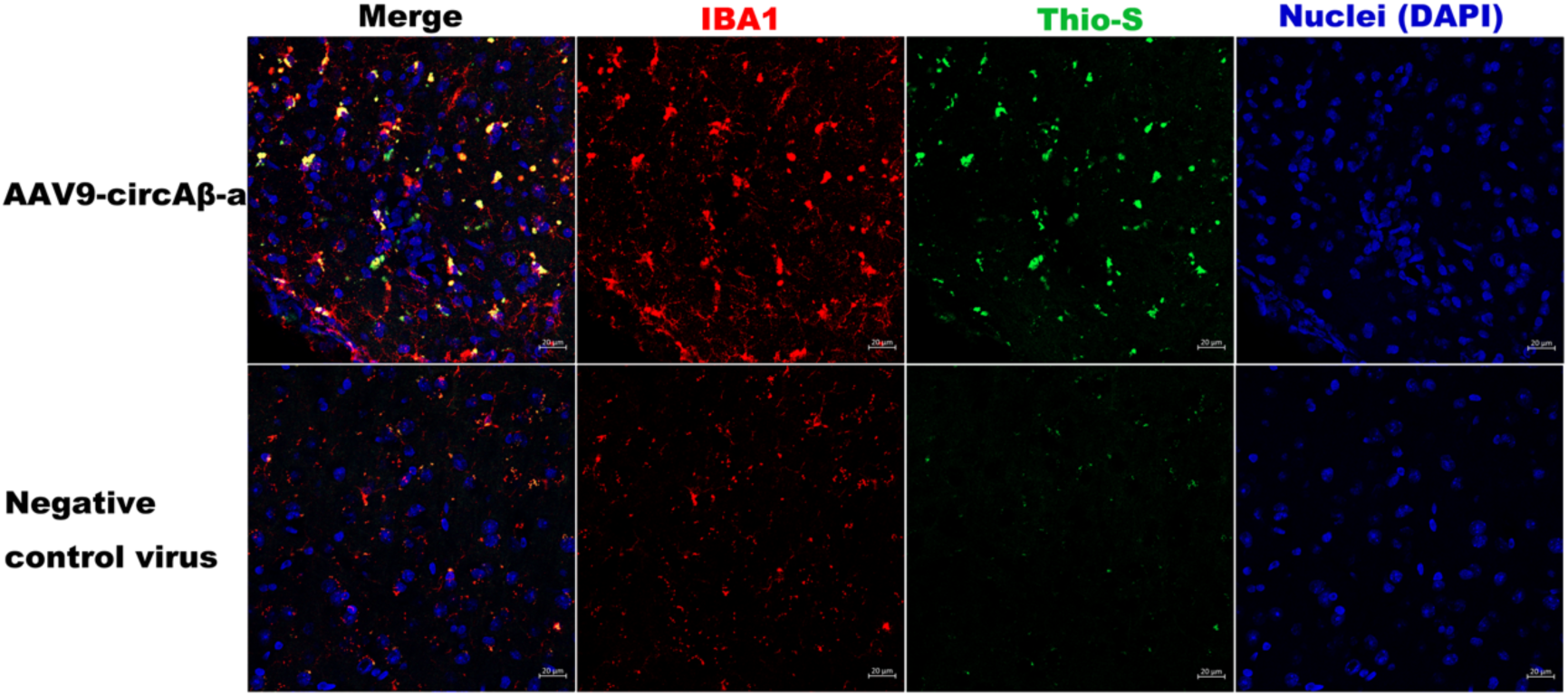
Microglial activation in the frontal cortex of AAV9-circAβ-a injected mice. blue, DAPI (nuclei); green, Thio-S; red, IBA1; Brain tissue from three mice were used (N=3); representative images were captured and shown here.

### Endogenous human Aβ175 expression in human HEK293 and HEK293T cell lines

The *in vivo* study of human circAβ-a expression in mouse brain demonstrates toxic Aβ peptide production as well as Aβ plaque formation. Whether such phenomena happen in human cells and especially human brains remains unknown. To overcome the low resolution and low sensitivity of Aβ antibodies in Western blots, we developed a novel antibody (MO-05) against the specific C-terminal region of Aβ175, which recognized the overexpressed Aβ175 protein (19.2 kDa) from the pCMV-Aβ175 construct in HEK293T cells with a band migration somewhat faster than 20 kDa in SDS-PAGE gels (Fig. 4A). Importantly, the antibody could also detect the endogenous Aβ175 in the negative control cells with empty vector transfection (Fig. 4A). For further verification of endogenous Aβ175 expression, we treated HEK293T cells with 300 nM specific antisense oligonucleotides (ASO) targeting the junction region of circAβ-a RNA, which encodes the unique C-terminal peptide that is not translated by the canonical linear APP mRNA.

**Fig. 4.**
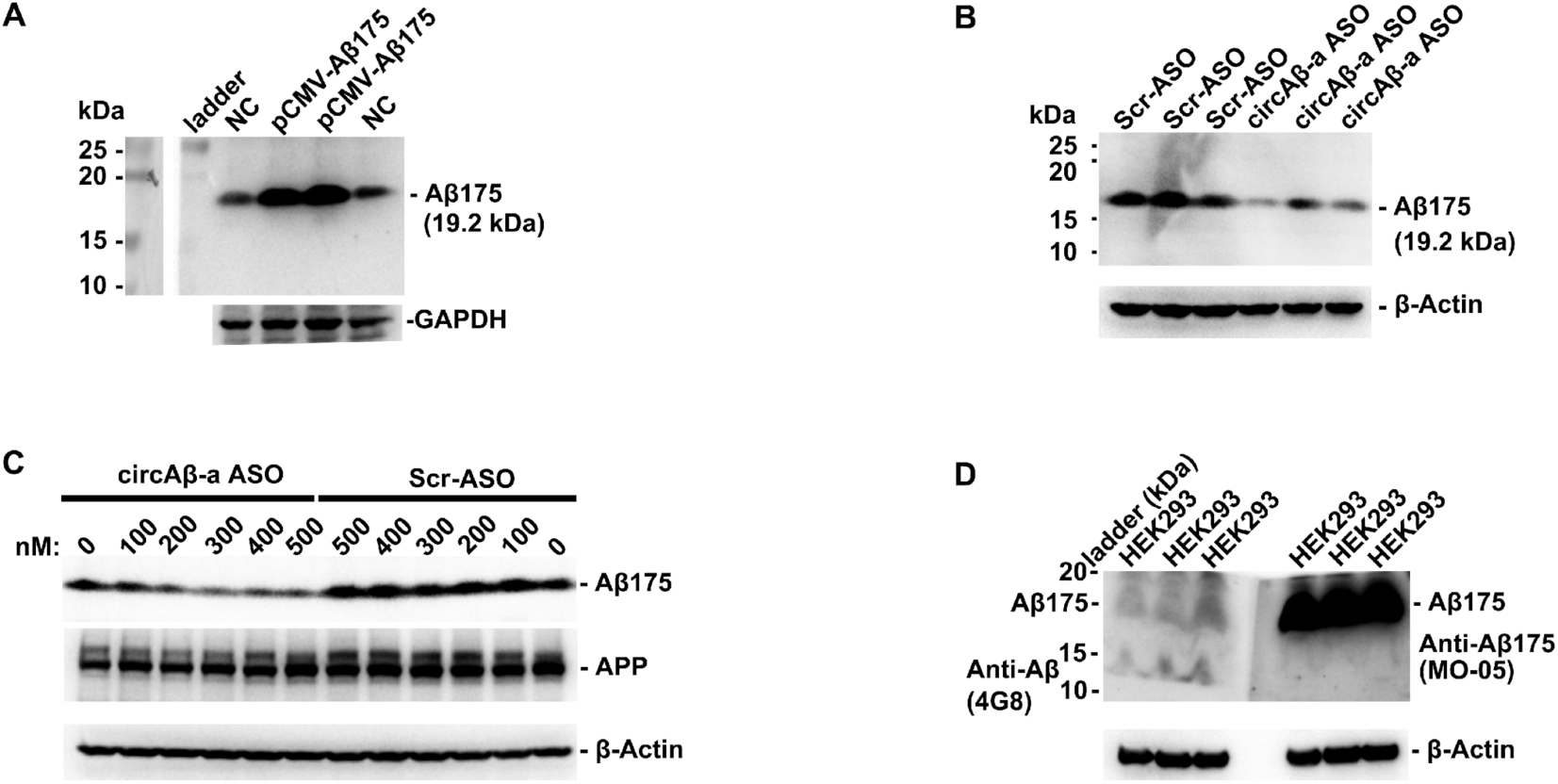
Endogenous Aβ175 expression in human HEK293T and HKE293 cell lines. **A**, overexpression of Aβ175 in human HEK293T cells. NC, empty vector transfection; pCMV-Aβ175 construct transfection. GAPDH was used as a loading control. Aβ175 was detected with MO-05 antibody (raised against the unique C-terminal sequence of Aβ175). **B**, ASO knock-down of Aβ175 expression in human HEK293T cells. Scr-ASO, 300 nM scrambled ASO transfection; circAβ-a ASO, the transfection of 300 nM ASO targeting the junction region of circAβ-a RNA. β-actin was used as loading control; Aβ175 was detected with MO-05 antibody. **C**, gradient concentration of ASO knock-down of Aβ175 expression in human HEK293 cells. Scr-ASO, scrambled ASO transfection; circAβ-a ASO, the transfection of ASO targeting the junction region of circAβ-a RNA. β-actin was used as loading control (A5441, Sigma); Aβ175 was detected with MO-05 antibody. **D**, detections of Aβ175 in three samples of untreated HEK293 cells by two different antibodies, respectively. The transferred membrane was divided into two parts. Left part was blotted with 4G8 (Aβ antibody); right part was blotted with MO-05 (Aβ175 antibody); the two parts were re-assembled and developed in ECL detection together.

Western blot analysis showed that compared to the scrambled ASO negative control, circAβ-a-ASO specifically and significantly downregulated endogenous Aβ175 levels (Fig.4B). Furthermore, we treated HEK293 cells with a range of ASO concentrations, where Aβ175 was reduced in the cells treated from 200 nM to 500 mM ASO, while the scrambled negative control ASO had no effect (Fig. 4C). Meanwhile, fl-APP protein levels remained the same in all these ASO-treated and untreated cells, demonstrating that our circAβ-a-ASO was highly specific (Fig. 4C).

One may wonder why previously Aβ175 had not been discovered in HEK293 or other cells using Aβ antibodies. We performed both 4G8 (Aβ antibody) and MO-05 (Aβ175 antibody) Western blots using the same membrane divided into two parts (Fig. 4D). The left membrane was incubated with 4G8 and the right membrane with MO-05 (Fig. 4D). When the two parts were juxtaposed, detection of the Aβ175 band was achieved with both antibodies and, at the identical position of the Aβ175 band detected with MO-05, we identified a faint band with 4G8 antibody (Fig. 4D). Despite the low sensitivity of 4G8 against Aβ175 (or fl-APP-derived CTFs) in Western blots, it further supported the similarity of the Aβ175 peptide with the Aβ related CTFs. The shortage of antibodies with sufficient sensitivity might explain why Aβ175 was overlooked in previous studies. When 4G8-incubated membranes were exposed for longer times, 4G8 did detect a strong band at the identical position of Aβ175 detected with MO-05 (Supplementary Fig. 2). Taken together, our results not only verified endogenous Aβ175 expression in human HEK293 cells, but also established the junction region of circAβ-a RNA as a viable target site for Aβ175 knock-down.

### Robust endogenous human Aβ175 expression in hNSC-derived human neurons, human brains and dysregulated Aβ175 pentamer processing during human brain ageing

As we have demonstrated the robust endogenous Aβ175 expression in human HEK293 and HEK293T cells, we asked whether Aβ175 is also expressed in human neurons and human brains. We generated human neuron cultures from human neural stem cells using a previously described protocol^32^. Western blot analysis showed that Aβ175 was robustly expressed in human NSC-derived neurons (Fig. 5A).

**Fig. 5.**
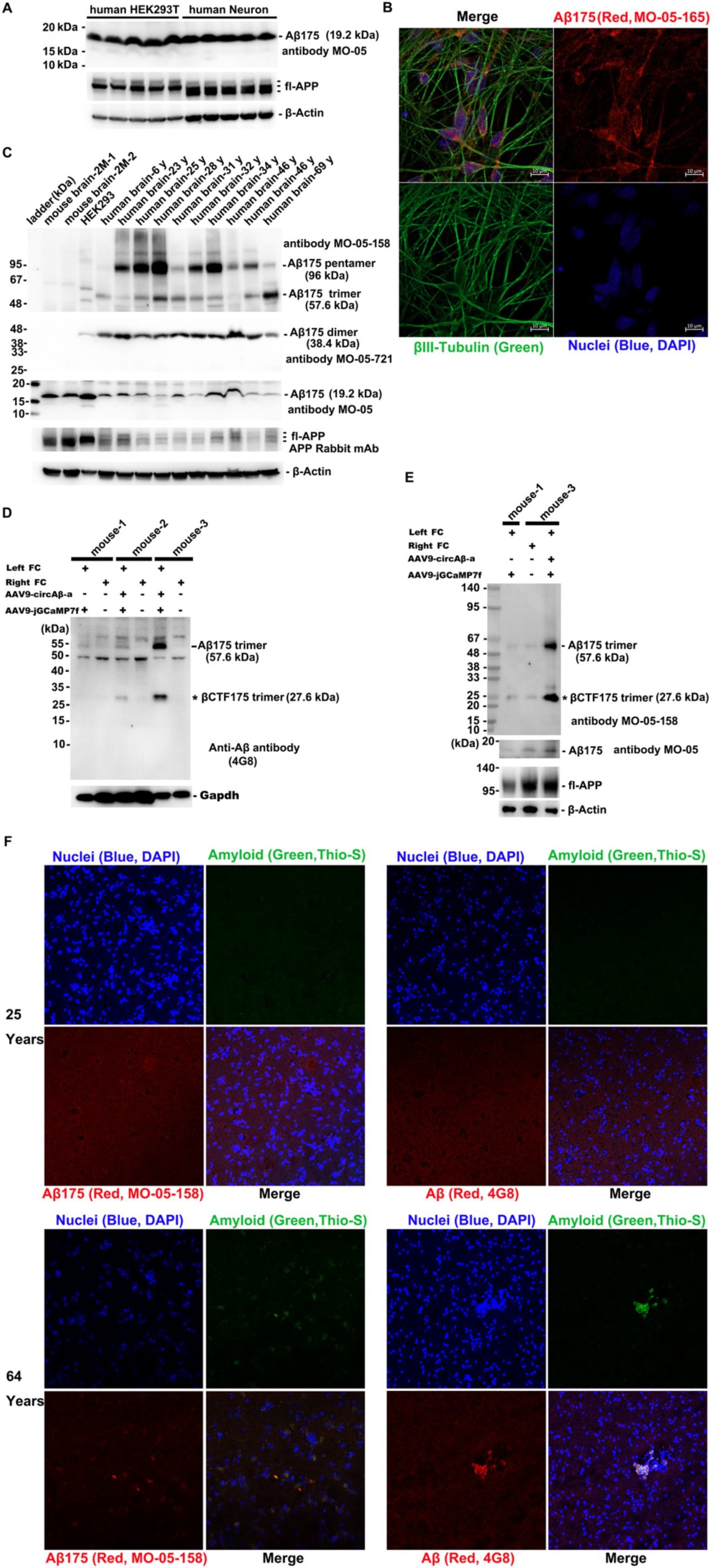
**Robust endogenous Aβ175 expression in a human neuronal cell line, human brains and dysregulated Aβ175 pentamer processing in the ageing brain.** A, Western blot analysis of the endogenous Aβ175 expression in hNSC-derived human neurons. β-Actin was used as loading control; fl-APP, full-length APP protein; five biological repeats were used here; B, Localization of Aβ175 in the soma of human neurons. Human neuronal culture was immunostained with anti-Aβ175 antibody (red; MO-05-165) and anti-βIII-tubulin rabbit mAb (green; A17913, ABclonal); Nuclei were stained with DAPI (blue). C, Western blot analysis of the Aβ175 oligomers in human and mouse brain total protein extracts (in RIPA buffer). Mouse brain-2M-1 and Mouse brain-2M-2 were brain frontal cortex samples from two-month-old C57BL/6 mice. Protein extracts from ten different human individuals were included (6–69 years of age, as indicated; for detailed sample information, see Supplementary Table 1). MO-05 also detected low levels of Aβ175 monomer expression in mouse brains (Fig. 5B). Divergent RT-PCR revealed that the mouse analog of circAβ-a RNA is expressed in mouse brains (Supplementary Fig. 5A, B). The mouse circAβ-a RNA also has an ORF which encodes Aβ175 similar in sequence to human Aβ175 (Supplementary Fig. 5C), explaining why MO-05 could also detect mouse Aβ175. D, E. Western blot analysis of Aβ175 oligomer formation in total protein extracts (in RIPA buffer) of AAV-circAβ-a injected mouse brain. Left FC, left frontal cortex; Right FC, right frontal cortex; Aβ was detected with 4G8 antibody; -, without; +, with virus injection. Mouse-1 was a 13-months-old negative mouse four months after control GFP-virus injection. The left frontal cortex of mouse-2 was injected with 0.70 × 10^10^ GC AAV-circAβ-a virus and mouse-3 with 6.26 × 10^10^ GC AAV-circAβ-a virus. Mouse Gapdh protein was used as loading control in D. β-Actin was used as loading control in E. *, putative βCTF175 trimer (27.6 kDa). As a note, Hepes-Tris-SDS-gel was used in D and MOPS-SDS-gel was used. E.F. Immunostaining of Aβ175 oligomers in young (25 years) and old (64 years) human brains. Nuclei, blue, DAPI staining; amyloid, green, Thioflavin S (Thio-S) staining; Aβ, red, 4G8 staining; Aβ175, red, MO-05-158 staining; yellow, merged red and green. Multiple slice images were captured, and representative images are shown. Twenty-five-year and 64-year-old human brain cortices were immunostained. Confocal microscopy (× 20) was used.

Immunofluorescence staining revealed that Aβ175 was mainly located in the soma of human neurons using monoclonal antibody MO-05-165 (described in Materials and Methods) against Aβ175 (Fig. 5B). We also detected Aβ175 in neurites (Fig. 5B). Our *ex vivo* results indicated that Aβ175 might be highly expressed in human neurons *in vivo* as well. Western blot analysis showed that Aβ175 monomer had only low-level expression in all the tested human brain samples from a large age range (6–69) (Fig. 5B). This was unexpected, as Aβ175 was highly expressed in *ex vivo* human neuron cultures.

We further wished to explore the presence of Aβ175 oligomers in human brains, as Aβ-related peptides are prone to oligomerization. Because the MO-05 antibody is polyclonal, and it can only detect Aβ175 monomer and dimer (data not shown), we generated several monoclonal antibodies which recognized Aβ175 oligomers, using the unique C-terminal peptide of Aβ175 as antigen (Supplementary Fig. 3). In human brain tissue, monoclonal antibody MO-05-158 could detect Aβ175 oligomers, including dimers, trimers, pentamers, heptamers and higher multimers, but not monomers. (Supplementary Fig. 3A). In particular, MO-05-158 detected high levels of Aβ175 pentamer formation in human brain (25-year-old) (Supplementary Fig. 3A). These results were confirmed by two other monoclonal antibodies (MO-05-160 and MO-05-165) (Supplementary Fig. 3B, C), highlighting the authenticity of Aβ175 pentamer. Furthermore, we used 4G8 and MO-05-158 to co-detect the same bands on Western blots which were cut into two parts (Supplementary Fig. 3E). The band corresponding to the Aβ175 pentamer in the young human brain samples detected by MO-05-158 could also be detected by 4G8, although at low sensitivity (Supplementary Fig. 3E). Furthermore, a longer SDS-gel run of total human protein extracts (pre-cleared over protein G beads) once again confirmed the detection of Aβ175 pentamer band by both MO-05-158 and 4G8 antibodies (Supplementary Fig. 3F).

Monoclonal antibody MO-05-721 could specifically detect Aβ175 dimers in human brain and human neuronal cultures, but not monomers and other oligomers (Supplementary Fig. 3D). shRNA lentivirus constructs, which specifically knocked down circAβ-a RNA significantly reduced Aβ175 dimer accumulation/formation in human neuronal cultures (Supplementary Fig. 3D). The formation of Aβ175 dimer in human brain was also confirmed by Aβ antibody 4G8 (Supplementary Fig. 3G).

With these powerful antibody tools, we detected robust Aβ175 oligomers in young adult human brain total protein extracts (Fig. 5C). As a note, MO-05-158 failed to detect any Aβ175 oligomer signals in two-month-old wild type mouse brains (mouse brain-2M-1 or 2 in Fig. 5C) as well. Compared to the human brains of three young adults in their twenties, which had robust Aβ175 pentamer accumulations, two human brains from individuals in their forties and one at age 69 exhibited remarkably lower levels of Aβ175 pentamers (Fig. 5C). In three brains from individuals in their thirties, Aβ175 pentamer levels declined in two cases and one sample maintained high levels (Fig. 5C).

We demonstrated that high levels of Aβ175 pentamers were present in the brains of young adult humans (twenties). The reduction of Aβ175 pentamers with age indicated that most Aβ175 pentamers derived from middle aged and older brains might be processed to yield Aβ peptides or aggregated into RIPA buffer unresolved plaques.

MO-05-721 antibody detected Aβ175 dimer accumulation in all human brain samples, with no significant changes across the age range (Fig. 5C). As a control, fl-APP expression levels remained the same in human brains of different ages (Fig. 5C); this is consistent with previous findings^37^.

We used MO-05-158, 4G8 and 1 µM thioflavin-S to co-stain brain slices derived from both young (25 years) and old (64 years) humans. We detected robust Aβ plaque signals with 1 µM thioflavin-S in old (64 years) but not young (25 years) brain slices and these plaques could be co-stained by 4G8 (Fig.5F), consistent with the previous findings that Aβ plaques appeared in the older individuals. MO-05-158 detected robust but diffuse signals in the young human brain (25 years), while the old brain sample (64 years) had significantly lower Aβ175 oligomer signals (Fig. 5F), consistent with Western blot analysis (Fig. 5C).

To investigate the oligomerization of Aβ175 in an *in vivo* mouse model, we performed Western blot analysis of AAV9-circAβ-a injected mouse brains with 4G8, MO-05 and MO-05-158 antibodies. As shown in Fig. 5D, E, mouse-3 exhibited a prominent signal in the left frontal cortex sample the of 6.26 × 10^10^ GC AAV9-circAβ-a virus injected sample at around 25 kDa. This band representing the putative human βCTF175 trimer (27.6 kDa) was detected by both 4G8 and MO-05-158 antibodies (Fig. 5D, E). As described above, in mouse-3 we detected strong Aβ expression, which further confirmed oligomerization of Aβ175. We also detected putative βCTF175 trimers with humanized MO-05-158 and MO-05-165 antibodies in young 23- and 25-year-old human brain samples (Supplementary Fig. 4A, B). We did not detect bands that would correspond to αCTF175 trimers (21.6 kDa) in either of the two human brain tissue samples, or the AAV9-circAβ-a virus injected brain tissue of mouse-3, (Fig. 5D, E, Supplementary Fig. 4), confirming that *in vivo*, Aβ175 trimers were predominately cleaved by β-secretase rather than α-secretase. There was no significant difference in Aβ175 monomer levels between GFP control mouse-1 and the circAβ-a expressing mouse-3 (Fig. 5E). The present results agree with our human brain data (Fig. 5C), where a major fraction of Aβ175 formed oligomers, and only a minor portion were detected as monomers.

Control mouse-1 did not have Aβ175 monomer and trimer formations detected by 4G8 (Fig. 5D). Failure to detect the endogenous mouse Aβ175 monomer in the control mouse brain presumably was caused by the low sensitivity of 4G8 antibody. The minor endogenous expression of mouse Aβ175 monomer and trimer in the 10.5-month-old mouse could be detected by our MO-05 and MO-05-158 antibodies (Fig. 5D, E). The mouse analog of circAβ-a RNA expressed in mouse brains was further described in Supplementary Fig. 5. Mouse-2 injected with a lower virus titer (0.70 × 10^10^ GC AAV9-circAβ-a virus) exhibited minor human Aβ175 trimer formation (Fig. 5D), while fl-APP protein levels remained unchanged between the left and right frontal cortex of mouse-3 (Fig. 5E), demonstrating that AAV9-circAβ-a injection did not affect endogenous mouse APP protein expression.

### Aβ175 the newly discovered component of Aβ plaques in older human brain tissue

Surprisingly, MO-05-158 also detected a robust signal in aggregates in the 64-year-old brain slices, and such protein aggregates were co-stained by thioflavin-S (the marker for amyloids) (Fig.5F), indicating that Aβ175 or its processing product(s) were component of Aβ plaques. To verify that the detected amyloid aggregates were, in fact, Aβ plaques, we then co-stained 64-year-old human brain slices with MO-05-158 (mouse mAb), thioflavin-S and β-Amyloid rabbit mAb (Fig. 6). Indeed, the MO-05-158 stained aggregates were co-stained by both thioflavin-S and β-Amyloid rabbit mAb, thus, confirming that these aggregates were Aβ plaques (Fig. 6). We concluded that aggregated Aβ175 oligomers were components of Aβ plaques in older human brain.

**Fig. 6.**
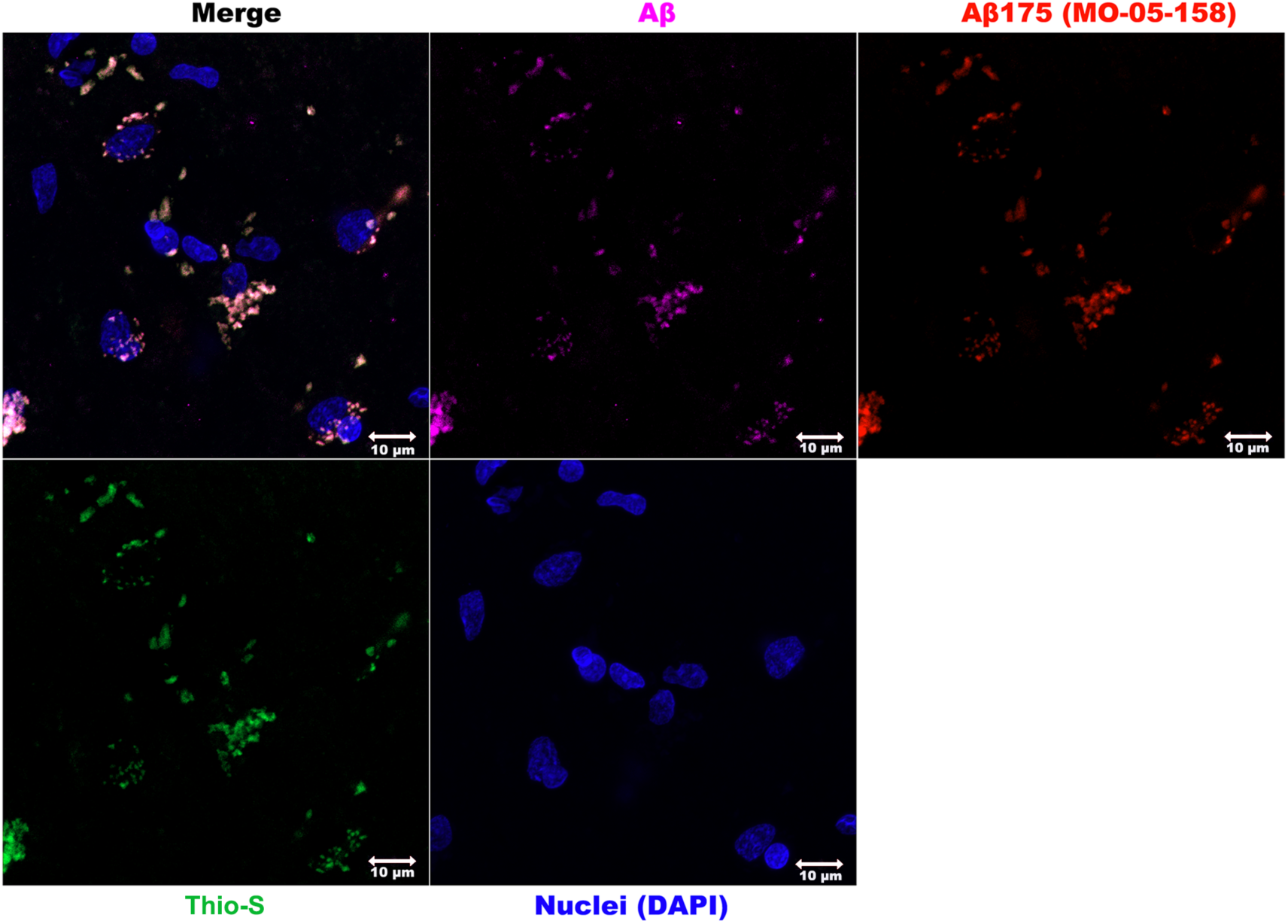
Co-staining of amyloids in brain slices of a 64-year-old human with Aβ175 oligomer antibody (red, MO-05-158), thioflavin-S (green) and β-Amyloid (1-40) rabbit mAb (purple). Nuclei were stained with DAPI (blue). Confocal microscopy (× 63) was used. Multiple images were captured with a representative shown here.

The inverted correlation between Aβ plaques and Aβ175 oligomers in young and old human brains strongly suggests that Aβ175 oligomers, especially Aβ175 pentamers, are processed to yield Aβ peptides or directedly aggregated into Aβ plaques, a process beginning in the mid-range age group (30s and 40s), further increasing with age and therefore should be considered a novel key driving force in the development of AD pathology. This would be consistent with previous findings of a strong correlation between Aβ plaque stages and ageing^38^.

We conclude that circAβ-a RNA was robustly translated into Aβ175 in hNSC-derived human neurons and human brains and propose that dysregulated proteolytic processing and aggregation of Aβ175 oligomers may cause Aβ plaque accumulation during human brain ageing.

### Aβ175, not fl-APP protein, produces the major Aβ peptides in HEK293 cells

As there are two potential precursors of Aβ peptides in human cells, it will be important to investigate which is the major source of Aβ. It is well accepted in the field that fl-APP protein is preferentially processed by α-secretase rather than β-secretase, which limits Aβ biogenesis^16^. Accordingly, Aβ175, instead of fl-APP, might be the major precursor of Aβ. To investigate this hypothesis, we used CRISPR-Cas9 genome editing technology to mutate exon-13 of the human APP gene in HKE293 cells to introduce a premature stop codon in endogenous fl-APP protein. The resulting APP truncation would not affect Aβ175 expression as the latter is encoded on exon-14 to exon-17 of the *APP* gene. We established five independent APP exon-13 edited cell lines. Western blots using KO validated APP rabbit mAb showed that fl-APP expression was abolished, while Aβ175 levels were not affected in these fl-APP-KO cell lines (Fig. 7A). We then measured the corresponding Aβ levels in the conditional medium of two fl-APP KO cell lines by indirect ELISA with 4G8 antibody. As shown in Fig. 7B, the two fl-APP KO cell lines still produced almost the same Aβ levels as the wild type HKE293 cells, demonstrating that KO of fl-APP protein had no detectable effect on Aβ expression. As fl-APP protein was absent in the fl-APP-KO cell lines, the detected Aβ could hardly be derived from fl-APP protein, demonstrating that the detected Aβ was processed from another precursor. Meanwhile, we used the shRNA lentivirus to specifically knock-down circAβ-a RNA expression in HEK293 cells, and then measured medium Aβ levels as described above. We found that the specific shRNA significantly reduced Aβ175 expression to 0.65-fold in wild type cells without affecting fl-APP expression levels (Fig. 7C, D). Aβ indirect ELISA demonstrated that extracellular Aβ was significantly reduced to 0.62-fold in the Aβ175 knock-down cells (Fig. 7D). The linear correlation between extracellular Aβ level and intracellular Aβ175 expression in HEK293 cells once more confirmed that Aβ175 is the major precursor of Aβ.

**Fig. 7.**
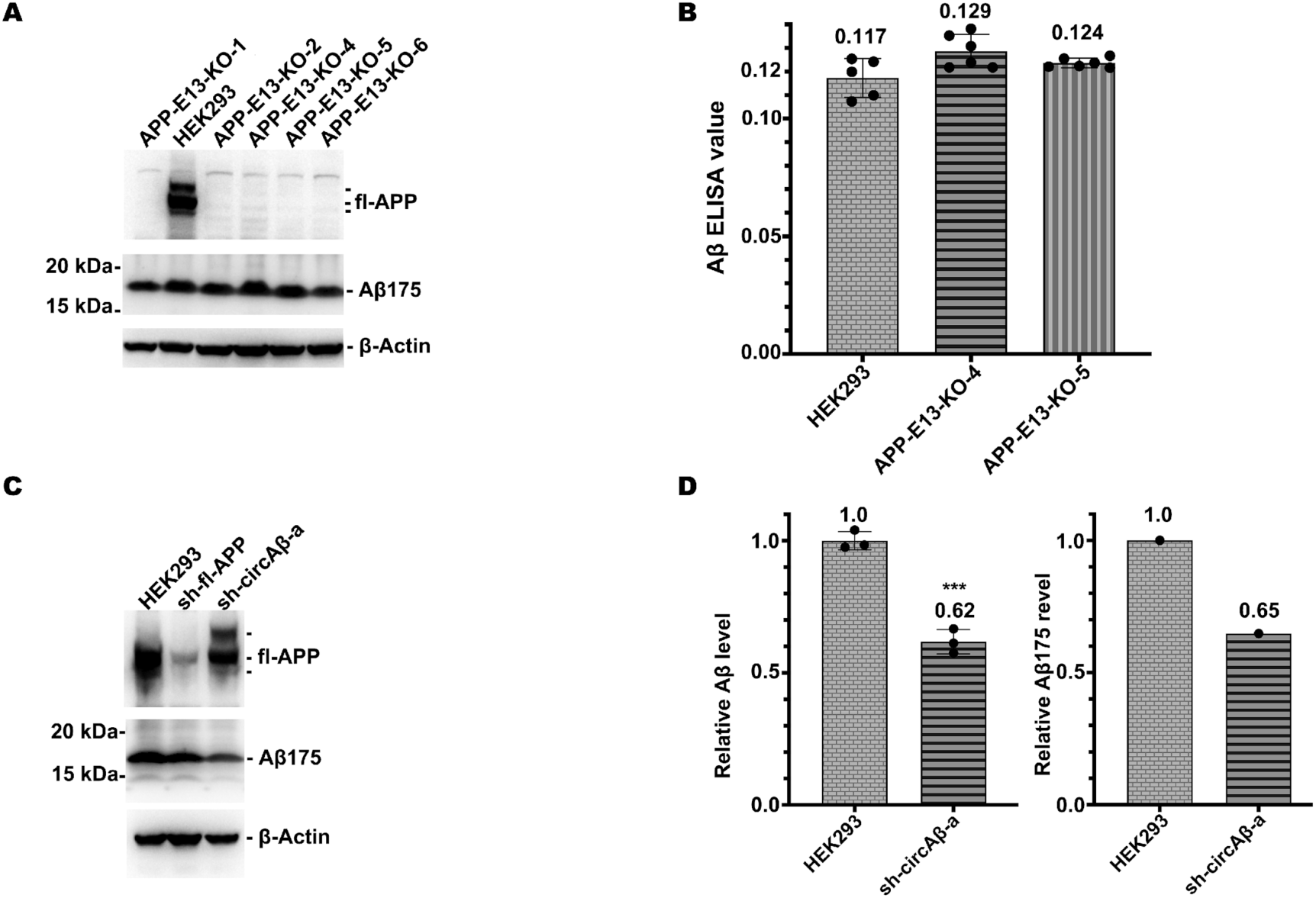
Aβ175, not fl-APP, is the major Aβ precursor in human HEK293 cells. A, Western blot analysis of fl-APP and Aβ175 expression in fl-APP-KO HEK293 cell lines. Aβ175 was detected with MO-05 antibody. fl-APP was detected by KO validated APP rabbit mAb (A17911, ABclonal). **B,** ELISA analysis of Aβ levels in the conditional medium of HEK293 wild type and fl-APP-KO HEK293 cell lines. **C,** Western blot analysis of fl-APP and Aβ175 expression in untreated and shRNA-lentivirus treated human HEK293 cells. HEK293, untreated HEK293 cells; sh-fl-APP, lentivirus treated HEK293 cells expressing shRNA targeting fl-APP mRNA; sh-circAβ-a, lentivirus treated HEK293 cells expressing shRNA targeting the junction of circAβ-a RNA; Aβ175 was detected with MO-05 antibody; fl-APP was detected by KO validated APP rabbit mAb (A17911, ABclonal). **D,** ELISA analysis of Aβ levels in the conditional medium of wild type and shRNA-lentivirus treated human HEK293 cells. The relative Aβ levels were calculated with reference to total protein. The relative Aβ175 levels were calculated with reference to β-Actin in C.

Taken together, fl-APP knockout and Aβ175 knock-down experiments solidly demonstrated that Aβ175, not fl-APP, produced the majority of Aβ peptides in human HEK293 cells, strongly indicating Aβ175 as the major Aβ precursor in human brains, especially in sporadic AD patients.

## Discussion

Alzheimer’s disease is a devastating neurodegenerative disorder that threatens our ageing society. Our understanding of AD has significantly advanced since the discovery of Aβ peptides and the derived Aβ hypothesis^1,9,39^, along with other hypotheses, such as the presenilin hypothesis in familial AD^40^. Recent advances of anti-amyloid drugs have proved that these Aβ antibodies could partially slow down disease progression^41^. Unfortunately, disease-halting or disease preventing drug development based on the Aβ hypothesis encounters challenges, reflecting the incompleteness of our current knowledge about AD molecular mechanisms^4,24,39,42,43^. Specifically, the main pathway for Aβ biogenesis in sporadic AD is still largely unknown and it has been assumed that the accumulation of Aβ peptides in sporadic AD brains is due to the imbalances in Aβ production or failure of Aβ clearing^4^. Such assumptions, however, fail to explain that non-mutated APP full-length proteins are not readily processed into toxic Aβ peptides^23^. Furthermore, neither APP nor γ-secretase proteins are mutated or upregulated in most sporadic AD patients. Wild type fl-APP protein is predominately processed by α-secretase, which generates, in conjunction with γ-secretase, the non-toxic P3 fragment rather than the toxic Aβ peptides^16^. These lines of evidence imply that there is a yet undiscovered alternative Aβ biogenesis pathway in sporadic AD brains.

We recently reported that human circAβ-a RNA generated from the human *APP* gene primary transcript expresses an Aβ containing polypeptide, Aβ175; this peptide is further processed to Aβ, providing an alternative route for Aβ production^23^. Using multiple lines of evidence, we here demonstrated robust Aβ formation in human circAβ-a RNA overexpressing mouse brains *in vivo*. The human circAβ-a expression could perfectly explain most Aβ features of human AD brains, including the extracellular Aβ plaque depositions and the microglial activation around the aggregated plaques (Fig. 3).

As the key hallmark of AD, robust Aβ plaque formations and depositions are observed in middle-aged (12–13 months) mouse brains after only a three or four-month-long expression of human circAβ-a RNA, further strengthening its pathogenetic role in AD. Interestingly, there is a correlation between the Aβ plaque burden sizes and the age of the mice and/or the expression period of the AAV-circAβ-a virus (Supplementary Fig. 1), consistent with the age-dependent development of Aβ plaque stages in humans^38^. Furthermore, we detected discernible endogenous expression of Aβ175 (human circAβ-a RNA translation product) in human HEK293 cells, hNSC-derived human neurons, and human brains (Fig. 5), thus supporting the potential significance of the alternative Aβ precursor in human AD. Interestingly, young human brains (in their twenties) exhibited robust soluble Aβ175 oligomer formation while the middle-aged and older human brains had much lower soluble Aβ175 oligomers levels, suggesting that reduction of Aβ175 oligomers is concomitant with processing to Aβ or they are directly forming insoluble plaques, thus principally causing the Aβ plaque accumulation in the ageing human brain. It is also possible that the decline of Aβ175 oligomers was caused by reduced formation or translation of circAβ-a RNA during ageing. A recent study found a decrease of circAβ-a (hs_circ_0007665) RNA in entorhinal cortex of a group of AD post mortem samples with high but not low or intermediate ABC scores^44^. Nevertheless, our data showed that despite a decline in Aβ175 oligomers, Aβ plaques still could accumulate with age. Our observations agree with previous findings that demonstrated a relatively early onset of Aβ plaque formation in human individuals during their thirties and forties that steadily increased with age^38,45^, while brains from younger humans have low Aβ levels^46^. Previous studies report a 20–30–year interval between the beginning of Aβ plaque development and the onset of dementia^46,47^, whereby sporadic AD (late-onset AD) symptoms commence at age 65 or older. Thus, the accumulation of Aβ peptides and the first deposition of Aβ plaques in brain should begin around ages 35–45. Moreover, there is emerging evidence that middle-aged (roughly between ages 40–59) human brains may change significantly, setting the stage for the development of dementia in older individuals^48^. Dysregulation of Aβ175 oligomer processing and/or aggregation during the ages of thirties and forties matches well with previous clinical observations. It will be of great importance to decipher whether the dysregulation of Aβ175 oligomer proteolytic processing and/or aggregation initiates the accelerating changes of the middle-aged human brain or *vice versa*. Furthermore, this study also confirmed that brain fl-APP protein levels remained unchanged as age increased, being consistent with previous findings that fl-APP expression levels are unaffected in both AD samples and non-demented controls^37^.

Taken together, multiple direct and indirect evidence strongly imply Aβ175 oligomers as a major Aβ precursor in human brain ageing and the dysregulation of Aβ175 oligomer processing and/or aggregation in middle-aged and older brains plays a critical role in Aβ accumulation in sporadic AD. Furthermore, in the singular older human brain sample we examined, Aβ175 oligomers were part of the componentry of Aβ plaques. Thus, Aβ175 oligomers not only supply the precursor of building units of Aβ plaques but also directly take part in the forming of Aβ plaques, underscoring their essential role in AD pathology.

Previous kinetic studies have shown that compared to other substrates such as neuregulin, affinity and catalytic efficiency of β-secretase is reduced toward fl-APP protein^49^. Here, we could demonstrate in fl-APP KO and Aβ175-shRNA knock-down experiments that Aβ175, not fl-APP, was the major precursor of Aβ peptides in human HEK293 cells (Fig. 7). Accordingly, in Western blots, we detected bands matching the size of βCTF175 trimers (β-secretase cleaved processing intermediate of Aβ175 oligomers), but not of αCTF175 trimers (α-secretase cleaved processing intermediate of Aβ175 oligomers) both in human circAβ-a RNA injected mouse brain and human brain tissue (Supplementary Figure 4). Taken together, this strongly indicates that similar mechanisms might underlie the development of Alzheimer’s disease in the human brain. We also discovered endogenous mouse circAβ-a RNA and mouse Aβ175 protein similar to their human counterpart. However, wild type mouse brain does not develop Aβ plaques and animals never exhibit AD pathology. A circAβ-a copy (circAltas ID: mma-APP_0013) is also expressed in the brain of rhesus macaque^50^. The predicted rhesus macaque Aβ175 copy has the same protein sequence as human Aβ175 (Supplementary Fig. 5C). It had been reported that rhesus monkeys aged 26 or older with severe memory impairments (SMI) had abundant Aβ plaques, neurofibrillary tangles (NFTs) and neuronal loss in their brains, thus resembling the three key AD pathological hallmarks in humans^51^. It is likely that the predicted circRNA encoded rhesus macaque Aβ175 polypeptide also plays a pathological role in the monkey brain during ageing.

We acknowledge the significance of previous studies on *APP*, *PSEN1*, *PSEN2* and other gene mutations in familial AD. Our study focused on the potential role of Aβ175 in sporadic AD and does not exclude the significant roles of *APP* (encoding the linear messenger RNAs), *PSEN1, PSEN2* and other mutations in familial AD. Since most *PSEN1* mutations lead to reduced production instead of increasing Aβ from βCTF (APP-C99), these findings were in conflict with the amyloid hypothesis in these familial AD patients^11^. Apparently, if Aβ175 or its processed intermediate, βCTF175 was the main substrate of γ-secretase, this dilemma might be solved. It will be interesting to investigate the production of Aβ from a Aβ175 monomer or oligomers with these mutated γ-secretases, which may resolve the controversy between the amyloid and presenilin hypothesis in familial AD^11,40^. In particular, our Aβ175 oligomer specific antibody (MO-05-158) will be a useful tool to obtain further answers regarding AD development. Moreover, the discovery of Aβ175 oligomers (and subsequent investigations) will complement and further develop the Aβ oligomer hypothesis, thus solving the long-standing mystery of Aβ oligomer formation in the human brain^52,53^.

Our study has some limitations, as we examined only two older human brain samples and no tissues from confirmed Alzheimer patients were available. Nevertheless, we might not have detected any unexpected differences. With the discovery of an alternative Aβ precursor, our attention should be shifted to the largely unknown molecular and/or environmental events near the bifurcation leading either to a path without symptoms or to a progression leading to AD. Further studies are required to decipher the mechanism of how Aβ175 oligomer processing and aggregation is dysregulated during human brain ageing, and especially during AD and why Aβ175 polypeptide but not fl-APP would be processed preferentially by β-secretase, thus causing sporadic AD. Our antibodies, mouse model and antisense strategies not only represent novel tools to undertake these studies, but they also hold promise as a basis for therapeutic interventions.

## Conclusions

Based on the data presented in this study and other evidence from previous research, we propose that Aβ175 is the key or major pathogenic factor in AD, as it is possibly not only the precursor of the majority of toxic Aβ peptides but also directly forms Aβ plaques in sporadic Alzheimer’s disease (Fig. 8). Our findings might significantly revise and enrich the Aβ hypothesis (Fig. 8). Specifically, these observations might have further elucidated our understanding of the origin of toxic Aβ forms which drive neural dysfunction in sporadic Alzheimer’s disease. The models described here will be invaluable to determine the factors responsible for increased Aβ expression in AD patients versus unaffected individuals. Finally, the tested antisense oligonucleotide targeting circAβ-a and the developed Aβ175 antibodies in this study have the potential to lead the way to next generation disease modifying drugs. Such novel weapons will be crucial to combat Alzheimer’s disease—one of the greatest health challenges of our time.

**Fig. 8.**
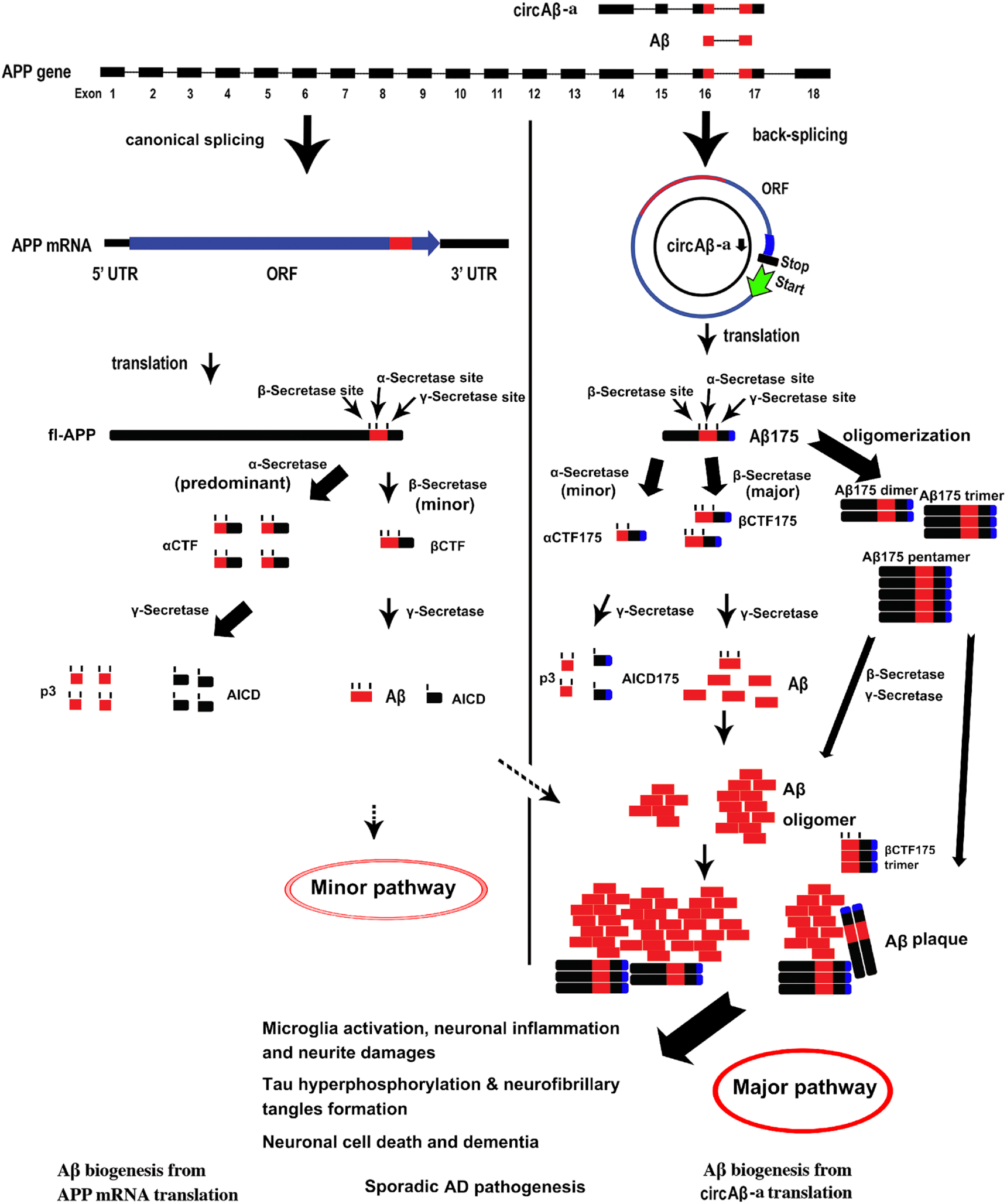
**circAβ RNA drives the formation of β-amyloid plaques in sporadic Alzheimer’s disease.**

Proposed mechanism of Aβ biogenesis and pathogenesis in sporadic Alzheimer’s disease. Modified from the previously published original model^23^. ORF, open reading frame; βCTF, fl-APP is cleaved by β-secretase to produce a C-terminal fragment (sometimes also called C99); βCTF175, circAβ-a encoded Aβ175 is cleaved by β-secretase to produce a C-terminal fragment; αCTF, fl-APP is cleaved by α-secretase to produce a C-terminal fragment and αCTF is further cleaved by γ-secretase to generate p3 and AICD; αCTF175, circAβ-a encoded Aβ175 is cleaved by α-secretase to produce a C-terminal fragment (this is hypothetical in this pathway, as we did not observe corresponding bands—neither monomer nor oligomer); AICD, amyloid precursor protein intracellular cytoplasmic/C-terminal domain, the smaller cleavage product of APP by γ-secretase; p3 peptide, also known as amyloid β-peptide (Aβ)_17– 40/42_; Aβ plaque, the amyloid plaque consisting of Aβ and Aβ175 (and possibly its further processing products.

## Ethics approval and consent to participate

All the animal experiments were performed according to the corresponding laws and regulations. Detailed animal protocol was approved by the IACUC committee at Peking University Shenzhen Graduate School. The using of human brain samples was reviewed and approved by the Ethic Committee of the Xiangya Hospital of Central South University. The research design and methods are in accordance with the requirements of regulations and procedures regarding to human subject protection laws such as GCP and ICH-GCP.

## Resource availability

### Lead contact

Requests for further information and resources should be directed to and will be fulfilled by the lead contact, Dingding Mo, School of Chemical Biology and Biotechnology, Peking University Shenzhen Graduate School, Shenzhen, 518055, China

### Materials and data availability

All the primary data and materials are available on request with a standard procedure to lead contract.

### Funding

G.X. was supported by National Natural Science Foundation of China (No. 82371362 and No. 82171347), Hunan Provincial Natural Science Foundation of China (No. 2022JJ30971), the Scientific Research Project of Hunan Provincial Health Commission of China (No. 202204040024). The development of Aβ175 antibodies and ASO used in the study were funded by D.M.

### Authors’ contributions

D.M. designed, conceived and supervised the study. D.M. performed experiments, analysed the results, prepared the figures, and wrote the manuscript. Y.Z. and Y.L. participated in data analysis. J.B. participated in data analysis, manuscript editing and revising. G.X. collected the human brain samples and participated in human brain data analysis.

## Acknowledgements

Lead author acknowledges Prof. Wen-Biao Gan, Dr. Wei Li, Cong Li, Dr. Hongling Guo and Baojun Zhang for support in the mouse model of this study. Lead author also acknowledges the Imaging Core Facility of Shenzhen Bay Laboratory and Shixian Huang for assistance in slides imaging. QM-FN-SO_3_ was a gift from Prof. Zhiqian Guo. Lead author acknowledges Dr. Bo Zhang for sharing regents during the study. Lead author acknowledges Dr. Sriram Balusu for sharing the Aβ plaque staining protocol. Authors also thank Stephanie Klco-Brosius for editorial advice.

## References

1 De Strooper, B. & Karran, E. The Cellular Phase of Alzheimer’s Disease. Cell 164, 603–615 (2016). 10.1016/j.cell.2015.12.056

2 Hyman, B. T. et al. National Institute on Aging-Alzheimer’s Association guidelines for the neuropathologic assessment of Alzheimer’s disease. Alzheimers Dement 8, 1–13 (2012). 10.1016/j.jalz.2011.10.007

3 Morris, G. P., Clark, I. A. & Vissel, B. Inconsistencies and controversies surrounding the amyloid hypothesis of Alzheimer’s disease. Acta Neuropathol Commun 2, 135 (2014). 10.1186/s40478-014-0135-5

4 Frisoni, G. B. et al. The probabilistic model of Alzheimer disease: the amyloid hypothesis revised. Nat Rev Neurosci 23, 53–66 (2022). 10.1038/s41583-021-00533-w

5 Hunter, S. & Brayne, C. Understanding the roles of mutations in the amyloid precursor protein in Alzheimer disease. Mol Psychiatry 23, 81–93 (2018). 10.1038/mp.2017.218

6 Selkoe, D. J. & Hardy, J. The amyloid hypothesis of Alzheimer’s disease at 25 years. EMBO Mol Med 8, 595–608 (2016). 10.15252/emmm.201606210

7 Zhang, Y., Chen, H., Li, R., Sterling, K. & Song, W. Amyloid beta-based therapy for Alzheimer’s disease: challenges, successes and future. Signal Transduct Target Ther 8, 248 (2023). 10.1038/s41392-023-01484-7

8 O’Brien, R. J. & Wong, P. C. Amyloid precursor protein processing and Alzheimer’s disease. Annu Rev Neurosci 34, 185–204 (2011). 10.1146/annurev-neuro-061010-113613

9 Hardy, J. The discovery of Alzheimer-causing mutations in the APP gene and the formulation of the "amyloid cascade hypothesis". FEBS J 284, 1040–1044 (2017). 10.1111/febs.14004

10 De Strooper, B. Loss-of-function presenilin mutations in Alzheimer disease. Talking Point on the role of presenilin mutations in Alzheimer disease. EMBO Rep 8, 141–146 (2007). 10.1038/sj.embor.7400897

11 Sun, L., Zhou, R., Yang, G. & Shi, Y. Analysis of 138 pathogenic mutations in presenilin-1 on the in vitro production of Abeta42 and Abeta40 peptides by gamma-secretase. Proc Natl Acad Sci U S A 114, E476–E485 (2017). 10.1073/pnas.1618657114

12 Bekris, L. M., Yu, C. E., Bird, T. D. & Tsuang, D. W. Genetics of Alzheimer disease. J Geriatr Psychiatry Neurol 23, 213–227 (2010). 10.1177/0891988710383571

13 Ahmed, R. R. et al. BACE1 and BACE2 enzymatic activities in Alzheimer’s disease. J Neurochem 112, 1045–1053 (2010). 10.1111/j.1471-4159.2009.06528.x

14 Kwart, D. et al. A Large Panel of Isogenic APP and PSEN1 Mutant Human iPSC Neurons Reveals Shared Endosomal Abnormalities Mediated by APP beta-CTFs, Not Abeta. Neuron 104, 256–270 e255 (2019). 10.1016/j.neuron.2019.07.010

15 Haass, C., Kaether, C., Thinakaran, G. & Sisodia, S. Trafficking and proteolytic processing of APP. Cold Spring Harb Perspect Med 2, a006270 (2012). 10.1101/cshperspect.a006270

16 Gabriele, R. M. C., Abel, E., Fox, N. C., Wray, S. & Arber, C. Knockdown of Amyloid Precursor Protein: Biological Consequences and Clinical Opportunities. Front Neurosci 16, 835645 (2022). 10.3389/fnins.2022.835645

17 Basi, G. S. et al. Amyloid precursor protein selective gamma-secretase inhibitors for treatment of Alzheimer’s disease. Alzheimers Res Ther 2, 36 (2010). 10.1186/alzrt60

18 Liu, C. X. & Chen, L. L. Circular RNAs: Characterization, cellular roles, and applications. Cell 185, 2016–2034 (2022). 10.1016/j.cell.2022.04.021

19 Mo, D. et al. A universal approach to investigate circRNA protein coding function. Sci Rep 9, 11684 (2019). 10.1038/s41598-019-48224-y

20 Wang, Y. et al. Expanding uncapped translation and emerging function of circular RNA in carcinomas and noncarcinomas. Mol Cancer 21, 13 (2022). 10.1186/s12943-021-01484-7

21 Fan, X., Yang, Y., Chen, C. & Wang, Z. Pervasive translation of circular RNAs driven by short IRES-like elements. Nat Commun 13, 3751 (2022). 10.1038/s41467-022-31327-y

22 Hwang, H. J. & Kim, Y. K. Molecular mechanisms of circular RNA translation. Exp Mol Med 56, 1272–1280 (2024). 10.1038/s12276-024-01220-3

23 Mo, D. et al. Circular RNA Encoded Amyloid Beta peptides-A Novel Putative Player in Alzheimer’s Disease. Cells 9, 2196 (2020). 10.3390/cells9102196

24 Granzotto, A., Vissel, B. & Sensi, S. L. Lost in translation: Inconvenient truths on the utility of mouse models in Alzheimer’s disease research. Elife 13 (2024). 10.7554/eLife.90633

25 Saito, T. et al. Single App knock-in mouse models of Alzheimer’s disease. Nat Neurosci 17, 661–663 (2014). 10.1038/nn.3697

26 Sasaguri, H. et al. APP mouse models for Alzheimer’s disease preclinical studies. EMBO J 36, 2473–2487 (2017). 10.15252/embj.201797397

27 Lalonde, R., Fukuchi, K. & Strazielle, C. APP transgenic mice for modelling behavioural and psychological symptoms of dementia (BPSD). Neurosci Biobehav Rev 36, 1357–1375 (2012). 10.1016/j.neubiorev.2012.02.011

28 Dourlen, P., Kilinc, D., Malmanche, N., Chapuis, J. & Lambert, J. C. The new genetic landscape of Alzheimer’s disease: from amyloid cascade to genetically driven synaptic failure hypothesis? Acta Neuropathol 138, 221–236 (2019). 10.1007/s00401-019-02004-0

29 Glazar, P., Papavasileiou, P. & Rajewsky, N. circBase: a database for circular RNAs. RNA 20, 1666–1670 (2014). 10.1261/rna.043687.113

30 Ran, F. A. et al. Genome engineering using the CRISPR-Cas9 system. Nat Protoc 8, 2281–2308 (2013). 10.1038/nprot.2013.143

31 Hong, C. S., Goins, W. F., Goss, J. R., Burton, E. A. & Glorioso, J. C. Herpes simplex virus RNAi and neprilysin gene transfer vectors reduce accumulation of Alzheimer’s disease-related amyloid-beta peptide in vivo. Gene Ther 13, 1068–1079 (2006). 10.1038/sj.gt.3302719

32 Li, W. et al. Rapid induction and long-term self-renewal of primitive neural precursors from human embryonic stem cells by small molecule inhibitors. Proc Natl Acad Sci U S A 108, 8299–8304 (2011). 10.1073/pnas.1014041108

33 Le, T. V., Crook, R., Hardy, J. & Dickson, D. W. Cotton wool plaques in non-familial late-onset Alzheimer disease. J Neuropathol Exp Neurol 60, 1051–1061 (2001). 10.1093/jnen/60.11.1051

34 Fu, W. et al. Rational Design of Near-Infrared Aggregation-Induced-Emission-Active Probes: In Situ Mapping of Amyloid-beta Plaques with Ultrasensitivity and High-Fidelity. J Am Chem Soc 141, 3171–3177 (2019). 10.1021/jacs.8b12820

35 Serrano-Pozo, A. et al. Stable size distribution of amyloid plaques over the course of Alzheimer disease. J Neuropathol Exp Neurol 71, 694–701 (2012). 10.1097/NEN.0b013e31825e77de

36 Liu, P. et al. Quantitative Comparison of Dense-Core Amyloid Plaque Accumulation in Amyloid-beta Protein Precursor Transgenic Mice. J Alzheimers Dis 56, 743–761 (2017). 10.3233/JAD-161027

37 Yang, L. B. et al. Elevated beta-secretase expression and enzymatic activity detected in sporadic Alzheimer disease. Nat Med 9, 3–4 (2003). 10.1038/nm0103-3

38 Braak, H. & Braak, E. Frequency of stages of Alzheimer-related lesions in different age categories. Neurobiol Aging 18, 351–357 (1997). 10.1016/s0197-4580(97)00056-0

39 Walsh, D. M. & Selkoe, D. J. Amyloid beta-protein and beyond: the path forward in Alzheimer’s disease. Curr Opin Neurobiol 61, 116–124 (2020). 10.1016/j.conb.2020.02.003

40 Shen, J. & Kelleher, R. J., 3rd. The presenilin hypothesis of Alzheimer’s disease: evidence for a loss-of-function pathogenic mechanism. Proc Natl Acad Sci U S A 104, 403–409 (2007). 10.1073/pnas.0608332104

41 Ramanan, V. K. & Day, G. S. Anti-amyloid therapies for Alzheimer disease: finally, good news for patients. Mol Neurodegener 18, 42 (2023). 10.1186/s13024-023-00637-0

42 Karran, E. & De Strooper, B. The amyloid hypothesis in Alzheimer disease: new insights from new therapeutics. Nat Rev Drug Discov 21, 306–318 (2022). 10.1038/s41573-022-00391-w

43 Benn, J. et al. Aggregate-selective removal of pathological tau by clustering-activated degraders. Science 385, 1009–1016 (2024). 10.1126/science.adp5186

44 Urdanoz-Casado, A. et al. circRNA from APP Gene Changes in Alzheimer’s Disease Human Brain. Int J Mol Sci 24 (2023). 10.3390/ijms24054308

45 Bischof, G. N., Rodrigue, K. M., Kennedy, K. M., Devous, M. D., Sr. & Park, D. C. Amyloid deposition in younger adults is linked to episodic memory performance. Neurology 87, 2562–2566 (2016). 10.1212/WNL.0000000000003425

46 Jansen, W. J. et al. Prevalence of cerebral amyloid pathology in persons without dementia: a meta-analysis. JAMA 313, 1924–1938 (2015). 10.1001/jama.2015.4668

47 Jia, J. et al. Biomarker Changes during 20 Years Preceding Alzheimer’s Disease. N Engl J Med 390, 712–722 (2024). 10.1056/NEJMoa2310168

48 Dohm-Hansen, S. et al. The ’middle-aging’ brain. Trends Neurosci 47, 259–272 (2024). 10.1016/j.tins.2024.02.001

49 Ben Halima, S., et al. Specific Inhibition of beta-Secretase Processing of the Alzheimer Disease Amyloid Precursor Protein. Cell Rep 14, 2127–2141 (2016). 10.1016/j.celrep.2016.01.076

50 Wu, W., Ji, P. & Zhao, F. CircAtlas: an integrated resource of one million highly accurate circular RNAs from 1070 vertebrate transcriptomes. Genome Biol 21, 101 (2020). 10.1186/s13059-020-02018-y

51 Li, Z. et al. Naturally occurring Alzheimer’s disease in rhesus monkeys. bioRxiv, 2022.2010.2020.513120 (2022). 10.1101/2022.10.20.513120

52 Benilova, I., Karran, E. & De Strooper, B. The toxic Abeta oligomer and Alzheimer’s disease: an emperor in need of clothes. Nat Neurosci 15, 349–357 (2012). 10.1038/nn.3028

53 Shankar, G. M. et al. Amyloid-beta protein dimers isolated directly from Alzheimer’s brains impair synaptic plasticity and memory. Nat Med 14, 837–842 (2008). 10.1038/nm1782

