## Supplementary figures and table for "circAβ-a RNA encoded Aβ175—the hidden driver of β-amyloid plaque formation and deposition in sporadic Alzheimer’s disease"

#### Supplementary Fig. 1

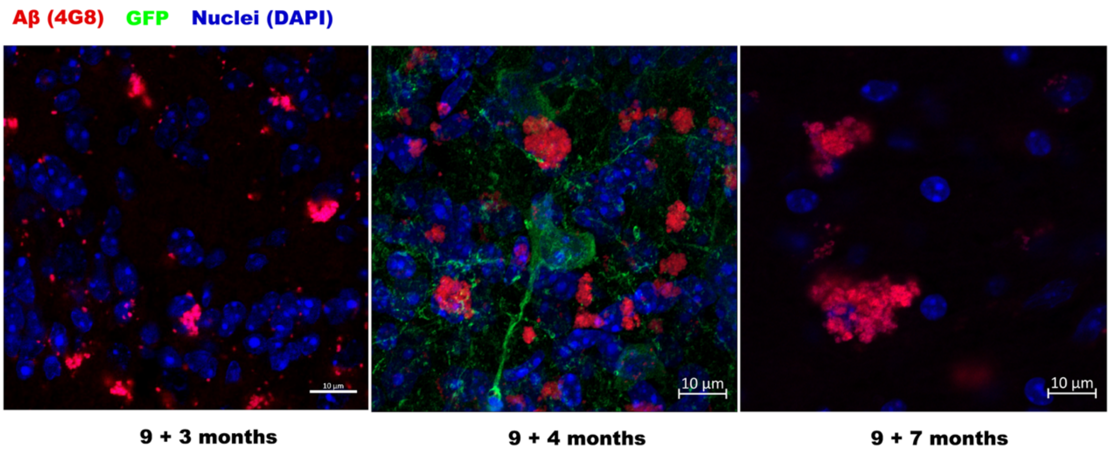


**Supplementary Fig. 1**

**Aβ plaque immunostaining of mouse frontal cortex injected with AAV9-SYN1-circAβ-a;** blue, DAPI (nuclei); red, Aβ (4G8); green, GFP; multiple slice images were captured; representative images are shown. 9+3 months, 9+4 months, 9+7 months corresponds to 3, 4, 7 months after injection of AAV9- circAβ-a into the frontal cortex of 9-month-old mice. Brain slices were collected and plaque immunostaining was performed with Aβ antibody (4G8). Control virus AAV9-jGCaMP7f expressing GFP was only injected into one mouse, namely the one examined after four months.

#### Supplementary Fig. 2

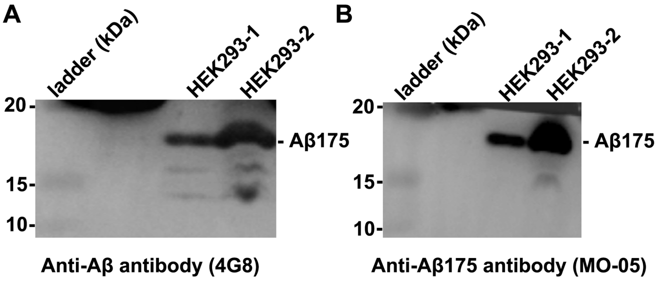

### Supplementary Fig. 2

**Western blot detection of Aβ175 by 4G8 and MO-05 antibodies in HEK293 cells.**

HEK293 cell total protein membrane was immunoblotted with 4G8 (Aβ antibody) and MO-05 (Aβ175 antibody); HEK293-1, HEK293-2, different amounts of HEK293 total cellular protein loaded.

### Supplementary Fig. 3

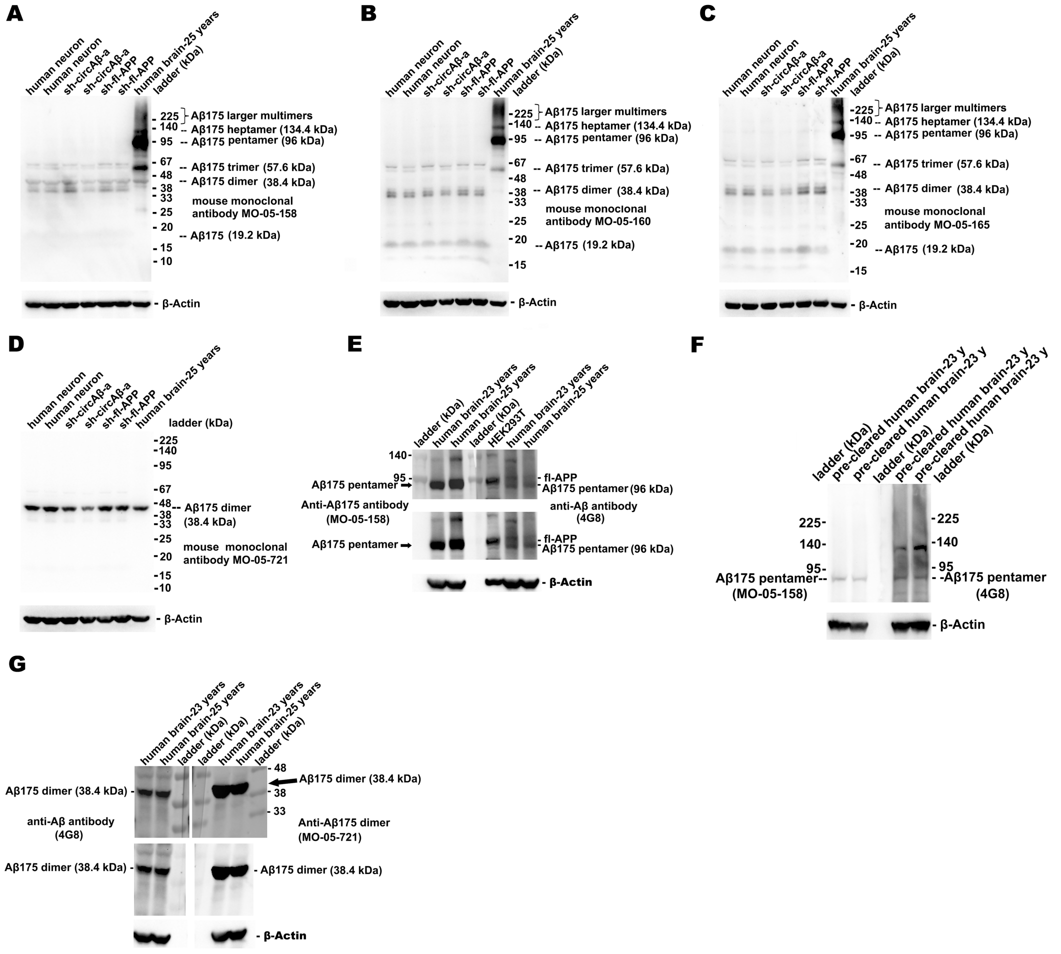


### Supplementary Fig. 3

**Specificity verification of Aβ175 oligomer antibodies.**

**A.** Aβ175 oligomers detection in human neuron culture and young human brains with monoclonal MO-05-158 antibody. β-Actin was used as loading control; human neuron, non-virus-infected human neuron culture; sh-circAβ-a, human neuron culture treated with sh-circAβ-a lentivirus (expression of shRNA targeting circAβ-a); sh-fl-

APP, human neuron culture treated with sh-circAβ-a lentivirus (expression of shRNA targeting APP mRNA); human brain-25 years, 25-year-old human brain sample.

**B, C.** Aβ175 monomer and oligomer detection in human neuron culture and young adult human brain with monoclonal MO-05-160, MO-05-165 antibody.

**D.** Aβ175 dimer detection in human neuron cultures and young adult human brain with monoclonal MO-05-721 antibody.

**E.** Double antibody detection of Aβ175 pentamers in young adult human brain. The transferred membrane was divided into two parts. The left part was blotted with MO-05-158 (Aβ175 oligomer antibody); the right part with 4G8 (Aβ antibody); the two parts were joined and co-developed in ECL.

**F.** Double antibody detection of Aβ175 dimers in young adult human brains. The transferred membrane was divided into two parts. The left part was blotted with 4G8 (Aβ antibody); the right part was blotted with MO-05-721 (Aβ175 oligomer antibody); the two parts were joined and co-developed in ECL.

**Supplementary Fig. 4**

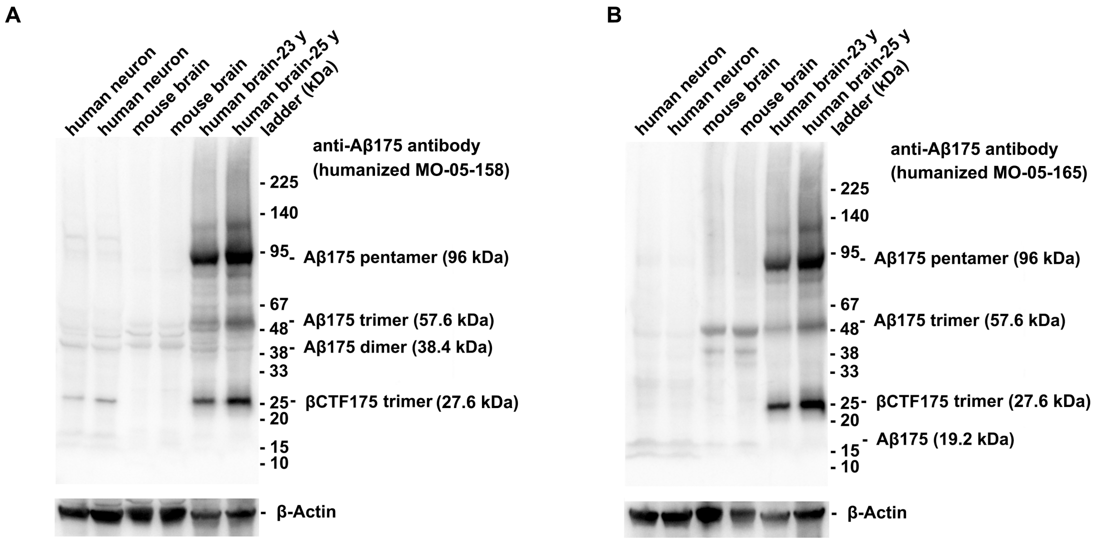


**Supplementary Fig. 4**

**Western blot detection of βCTF175 trimers and Aβ175 oligomers in human**

**brains.**

**A.** βCTF175 trimer and Aβ175 oligomer detection in young human brains (ages 23 and 25) with humanized MO-05-158 antibody. β-Actin was used as loading control; human neuron, non-virus-infected human neuron culture; mouse brain, un-virus-injected two-month-old mouse brain tissue; human brain-23 y, 23-year-old human brain sample; human brain-25 y, 25-year-old human brain sample. **B.** βCTF175 trimer and Aβ175 oligomer detection in young human brains with humanized MO-05-65 antibody.

### Supplementary Fig. 5

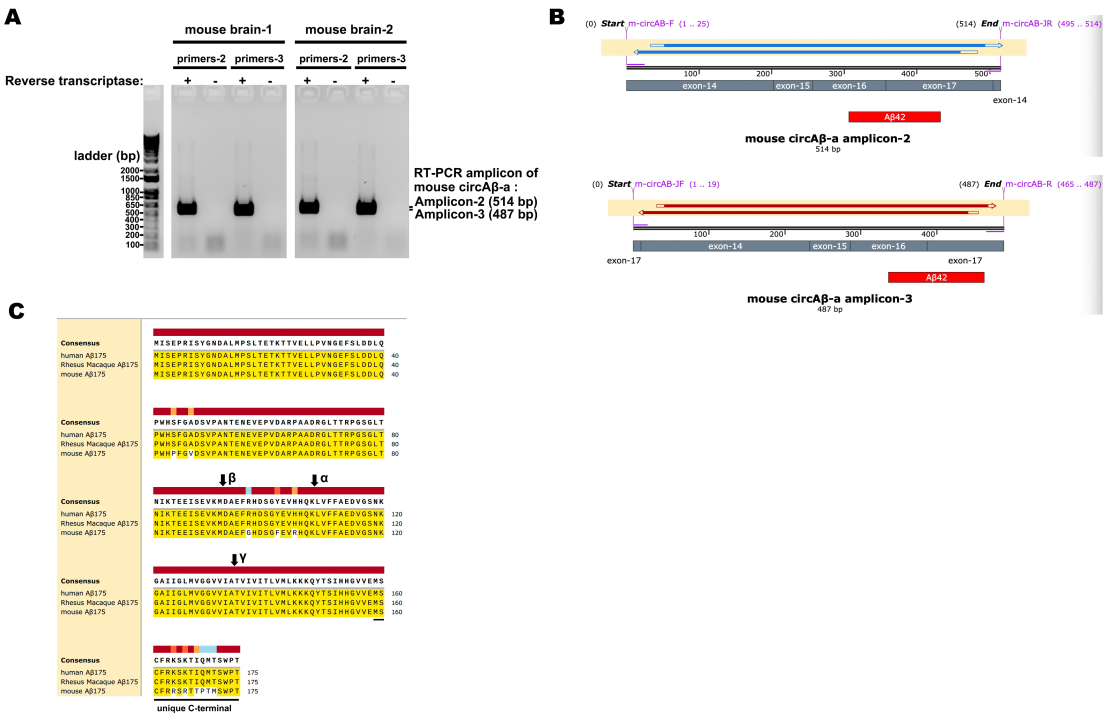


### Supplementary Fig. 5

**Mouse circAβ-a copy in mouse brain.**

**A.** RT-PCR of mouse circAβ-a copy in the total RNA of mouse brain; amplicon-2 with primer pair 2 was 514 bp (forward primer, 5’ GCGGCGTTGTCATAGCAACC 3’; reverse primer, 5’ GCACAGAGTCCACCCCAAAGG 3’); amplicon-3 with primer pair 3 was 487 bp (forward primer, 5’ CAACCGTGATTGTCATCACCCTGG 3’; reverse primer, 5’ CACGGCTGGAGGTCATCCAGG 3’);

**B.** Alignment of amplicon-2 and amplicon-3 sequencing.

**C.** Amino acid sequence alignment of human, rhesus macaque and mouse Aβ175 open reading frames using Clustal Omega. Consensus Threshold: >50%; rhesus macaque Aβ175 is the putative translation product from circAβ-a RNA copy (circAltas ID: mma-APP_0013). Amino acids that match the reference (human Aβ175) are marked with yellow highlighting. The unique C-terminal sequence is underlined. Vertical arrows indicate the predicted β-, α and γ-secretase cleavage sites in the circular RNA encoded Aβ175 polypeptides.

**Supplementary Table 1. Human brain tissue information**

| Patient Number | Sex | Age (years) | Sample of brain region | Disease | Cognitive ability | Used in experiments |
|---|---|---|---|---|---|---|
| 1 | NA | 6 | NA | NA | NA | Western blot |
| 2 | Male | 23 | Left frontotemporal lobe | Epilepsy | Epilepsy associated consciousness disorder | Western blot |
| 3 | Female | 25 | Temporal lobe | Cavernous hemangioma | Cognitive normal | Western blot |
| 4 | Female | 25 | Right frontotemporal lobe, hippocampus, basal ganglia | Glioma | Cognitive normal | Immunostaining |
| 5 | Male | 28 | Temporal lobe | Epilepsy | Epilepsy associated consciousness disorder | Western blot |
| 6 | Male | 31 | Left frontal lobe | Epilepsy | Epilepsy associated consciousness disorder | Western blot |
| 7 | Male | 32 | Temporal lobe | Cavernous hemangioma | Cognitive normal | Western blot |
| 8 | Male | 34 | Right hippocampus | Epilepsy | Epilepsy associated consciousness disorder | Western blot |
| 9 | Female | 46 | Left hippocampus | Epilepsy | Epilepsy associated | Western blot |

| | | | | | | |
|---|---|---|---|---|---|---|
| | | | | | consciousness disorder | |
| 10 | Male | 46 | Left temporal lobe | Epilepsy | Epilepsy associated consciousness disorder | Western blot |
| 11 | Female | 64 | Left occipital lobe | Glioma | Cognitive normal | Immunostaining |
| 12 | Female | 69 | Left lateral ventricle | Hematoma | Cognitive normal | Western blot |

NA, not available.
